# Ligand-induced shifts in conformational ensembles that predict transcriptional activation

**DOI:** 10.1101/2022.05.29.493907

**Authors:** Sabab Hasan Khan, Sean M Braet, Stephen John Koehler, Elizabeth Elacqua, Ganesh S Anand, C. Denise Okafor

## Abstract

Nuclear receptors function as ligand-regulated transcription factors whose ability to regulate diverse physiological processes is closely linked with conformational changes induced upon ligand binding. Understanding how conformational populations of nuclear receptors are shifted by various ligands could illuminate strategies for the design of synthetic modulators to regulate specific transcriptional programs. Here, we investigate ligand-induced conformational changes using a reconstructed, ancestral nuclear receptor. By making substitutions at a key position, we engineer receptor variants with altered ligand specificities. We use atomistic molecular dynamics (MD) simulations with enhanced sampling to generate ensembles of wildtype and engineered receptors in combination with multiple ligands, followed by conformational analysis and prediction of ligand activity. We combine cellular and biophysical experiments to allow correlation of MD-based predictions with functional ligand profiles, as well as elucidation of mechanisms underlying altered transcription in receptor variants. We determine that conformational ensembles accurately predict ligand responses based on observed population shifts, even within engineered receptors that were constitutively active or transcriptionally unresponsive in experiments. These studies provide a platform which will allow structural characterization of physiologically-relevant conformational ensembles, as well as provide the ability to design and predict transcriptional responses in novel ligands.

## Introduction

Nuclear receptors (NRs) are master regulators of diverse physiological functions, including reproduction, inflammation, development and metabolism [1-7]. The ability of this class of ligand-activated transcription factors to control critical cellular functions is driven by binding of lipophilic ligands. Members of the NR superfamily include steroid hormone receptors, such as the well-known estrogen and progesterone receptors, which are regulated by cholesterol-derived hormones. NRs share a characteristic modular domain architecture, including a highly conserved DNA binding domain and a moderately conserved ligand binding domain (LBD) which houses a hydrophobic binding cavity. In steroid receptors, ligand binding induces a conformational change in the receptor, followed by dissociation from chaperone proteins [8], binding to the DNA response elements located in promoter regions of target genes [9-12], and recruitment of coregulator proteins [13-15].

Differential conformational changes induced by ligands permit the recruitment of coregulator proteins that either promote transcriptional activation, or repression of target genes [14, 16, 17]. Activating ligands (agonists) will switch the receptor into a so-called active state, generally defined by the conformation in which the C-terminal helix (i.e. helix 12) is packed against the LBD, stabilized by interactions with nearby helices 3 and 4, as well as ligand contacts [18-21]. This region comprises the activation function 2 (AF-2) surface [22]. The active state positioning of H12 allows coactivator proteins to be recruited to the LBD [23-25]. Conversely, repositioning or destabilization of H12 is associated with inactive states of receptors such as an apo state or an antagonist-bound state [18, 26]. However, experimental evidence indicates that NRs exist as a dynamic ensemble of conformations whose populations can be modulated by ligand binding or other perturbations [27]. While it is an ongoing challenge to structurally characterize NR conformational ensembles and reveal ligand-induced population shifts, experimental methods such as solution state NMR have enabled great advances, revealing how ligands of diverse efficacy and potency affect the active state [28, 29].

Powerful advancements in computational approaches have increased their application for the study of protein conformational ensembles. Computational methods for conformational sampling are notoriously hampered by two major challenges: inherent limitations in forcefields, and the difficulty of achieving sufficient sampling of the free energy landscape [30, 31]. Enhanced sampling methods applied to molecular dynamics (MD) simulations, including accelerated MD, metadynamics and replica exchange have been useful for overcoming limitations in studying conformational ensembles, providing physical descriptions that illuminate structural and functional protein mechanisms [32-37]. Previously we showed that conformational ensembles of NRs generated by accelerated MD simulations underwent conformational shifts upon addition of ligands [38]. Unexpectedly, the population shifts correlated with the transcriptional activity of the ligands. Thus, understanding the effects of ligands on NR ensembles can be a promising approach for screening and predicting functional profiles of new NR ligands.

In this work, we expand our previous system by characterizing conformational shifts in a set of engineered receptors with altered ligand specificity and transactivation potential. We investigate conformational ensembles using a reconstructed ancestral steroid receptor, AncSR2. In transcriptional assays, AncSR2 was activated by 3-ketosteroid hormones, i.e. steroids with a non-aromatized A-ring and a keto substituent at the carbon 3 position, while remaining unresponsive to hormones with an aromatic A-ring, i.e. estrogens (**Fig. 1A**) [39]. To produce a diverse set of receptors with a range of functional profiles, we created four AncSR2 variants by mutating M75, a critical residue located on helix 5 (H5) of the LBD that is conserved across modern steroid receptors and shown by us and others [38, 40] to be crucial for hormone recognition. We use MD simulations to predict the conformational effects of M75 substitutions, generate conformational ensembles and predict population shifts that occur upon binding to aromatized and non-aromatized hormones. We then combine cellular assays with biophysical and structural analyses to dissect the structure-activity criteria underlying functional responses of AncSR2 variants to diverse ligands. Finally, we correlate experimental results with conformational shifts in computational ensembles to elucidate ligand-induced effects in a set of receptors.

**Figure 1:**
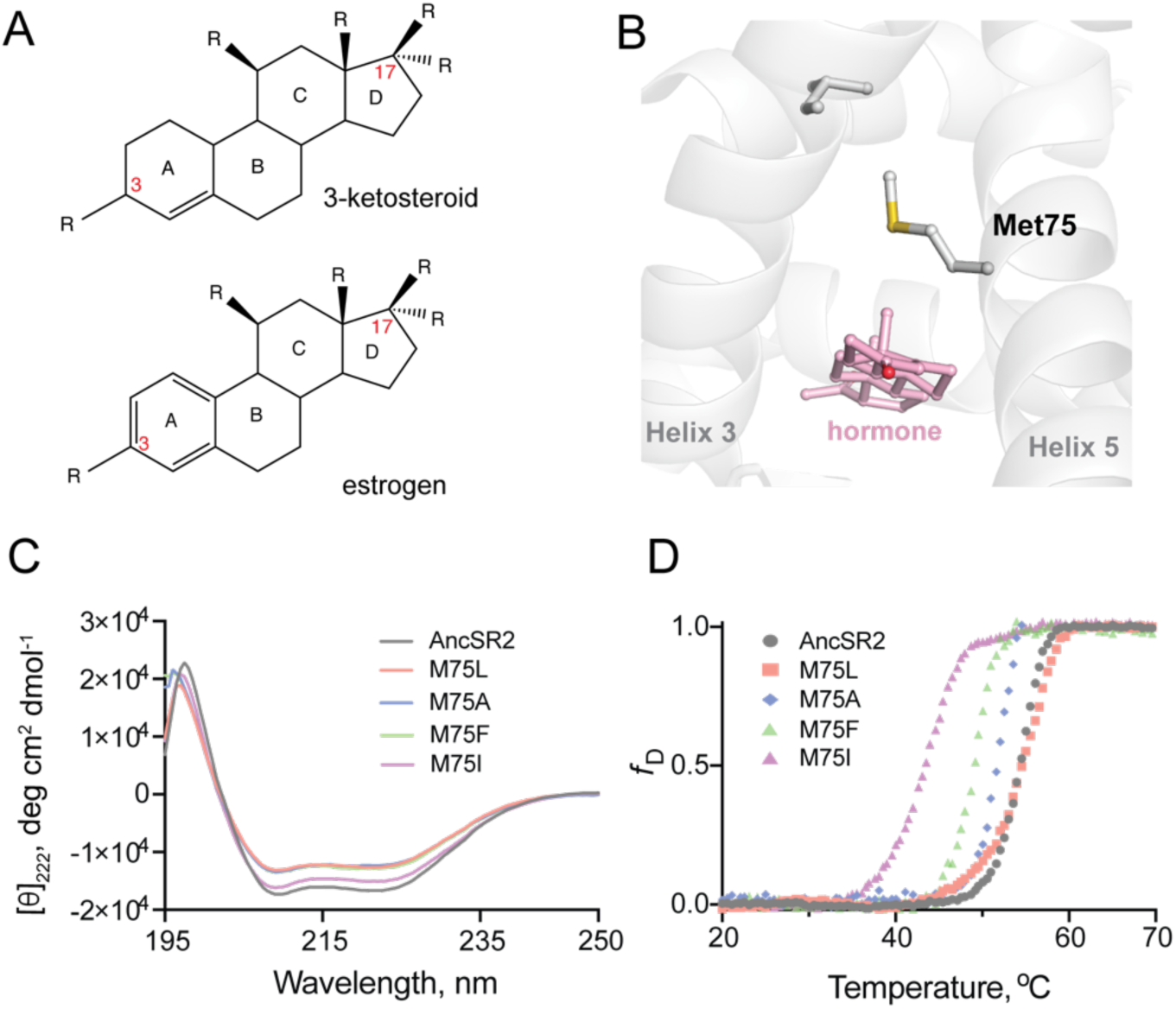
Structure and stability of AncSR2 and M75 mutants. **A)** Chemical structure of A-ring aromatized estrogen and non-aromatized 3-ketosteroid. **B)** Met75 (H5) is positioned to form critical contacts with H3 and the hormone located in the ligand binding pocket. **C)** Far-UV CD spectra of AncSR2 and its variants in the wavelength range 195-250 nm, showed mutants LBD remains in the folded state similarly as AncSR2. **D)** Normalized heat-induced denaturation transition curves of AncSR2 and its variants monitored by change in the [*θ*]_222_ as function of temperature. Each curve represents averaged measurements from two replicates and two independent purifications.

We observed that the M75 mutations achieved a striking range of functional profiles in AncSR2, including constitutively active and completely inactive states, enabling a broad investigation into how receptor function affects population shifts within NR ensembles. Our studies reveal that ensembles generated are extremely sensitive to both M75 mutations and ligand identity. In AncSR2 receptor variants, population shifts, assessed by clustering ligand-bound conformations with unliganded ensembles, correlated with functional properties of ligands. Changes in ensemble populations were predictive of strong, weak or no agonist activity in ligands. To reveal the origin of inactivity or constitutive activity in two of our variants, we employed hydrogen deuterium exchange mass spectrometry and ligand binding assays. We determined that shifts within conformational populations also accurately predicted the ligand response to these unexpected variants, confirming the inherent promise in this approach for characterizing diverse NR-ligand ensembles.

## Results

### Substitution of M75 has minor effects on global structure and larger impact on local interactions

Met75, located on helix 5 (H5) of AncSR2 holds structural, functional, and evolutionary significance for steroid receptors. Notably, M75 engages H3 residues via van der Waals contact **(Fig. 1B)**, an interaction that is conserved in extant glucocorticoid, mineralocorticoid, and progesterone receptors [41]. M75 was also shown to interact with bound hormones [40] representing an ideal position for mutagenesis to create a series of engineered receptors with altered potency and ligand specificity. We generated M75L, M75I, M75F, M75A mutants of AncSR2 and performed biophysical characterizations to ensure that mutations do not substantially impact structure and stability. Wildtype (WT) AncSR2 LBD and mutants were expressed and purified to homogeneity **(Fig. S1)**. Gel filtration profiles show that similar to WT, mutant LBDs elute as a single peak, suggesting that mutations do not affect the globular nature of the protein (Data not shown). We used circular dichroism spectroscopy (CD) to determine the impact of M75 mutations on the structure of AncSR2. Far-UV CD (195-250 nm) spectral measurements of the WT AncSR2 and mutants reveal features characteristic of α-helical proteins, i.e., negative minima at 208 nm and 222 nm and positive maximum around 190 nm **(Fig. 1C)**. However, a slight decrease in the mean residue ellipticity at both negative minima was observed in mutants (M75L, M75A, M75F) as compared to WT AncSR2. Overall, CD measurements confirm that mutations do not affect the global secondary structure of AncSR2.

We also tested the effect of mutations on the stability of AncSR2 and M75 variants by following changes in the CD signal at 222 nm as a function of temperature (**Fig. 1D)**. The equilibrium denaturation curves for each protein were analyzed to obtain melting temperatures (*T*_m_). The apparent *T*_m_ values for M75L mutant and WT are identical, within experimental error. The apparent *T*_m_ values of M75A, M75F and M75I are respectively 2.3, 5.2 and 10.9 °C lower than AncSR2. Thus while mutant receptors retain secondary structural characteristics of WT AncSR2, stability is reduced in a few variants which may be reflective of local, structural effects.

From our previous work, it is known that contact between H3-H5, as well as interactions between M75 and hormones may predict how well a hormone can activate AncSR2 [38]. To reveal the impact of M75 mutations on residue and ligand contacts, we used MD simulations to model each variant in the presence of five hormones: four 3-ketosteroids (progesterone, DHT, hydrocortisone, aldosterone) and estradiol. We included an unliganded (apo) simulation for each variant, generating a total of thirty complexes. To measure contacts between two residues, we determined the minimum distance between heavy atoms (See Methods) of both residues across the simulations. First, we measured the distance between residue 75 and the hormone (**Fig. 2A)**. In all mutants and for all ligands, residue 75 is within 4.5 Å of the hormone which is within the threshold for a van der Waals contact (**Fig. 2B)**. Of all variants, M75I has the shortest distances for this contact, with values ranging from 3.5 -3.7Å. Distances in all other variants are higher, ranging from 3.7 - 4.0 Å.

**Figure 2:**
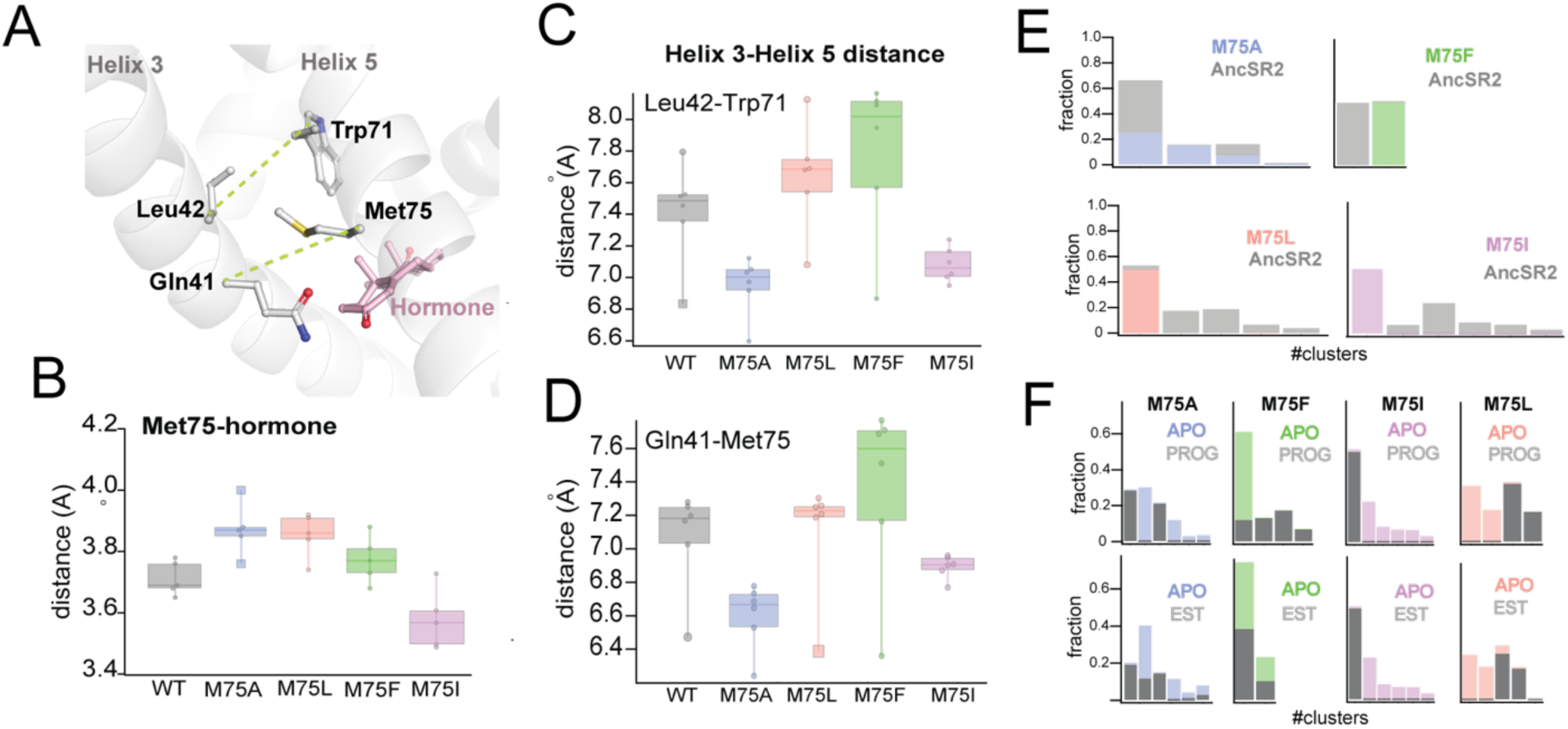
Contact measurements and clustering analysis from MD simulations. **A)** AncSR2 structure indicating positions of M75 along with bound hormone and additional H5 and H3 residues used to determine contact measurements. To assess H3-H5 interhelical distances, measurements were performed between Leu42-Trp71 and Gln41-res75. **B)** Average distances between residue 75 (H5) and A-ring of hormones. **C)** Average distances between L42 Cα and W71 Cα atoms. M75A showed the smallest distances, M75F the largest, with all other complexes in between, indicating that the size of the sidechain determines the interhelical distance. **D)** Average distances between Q41 Cα and res75 Cα atoms. This contact follows the same trends observed in **(C)**. Individual data points in B, C, D represent distance measurements averaged over simulations. Each box and whisker representation displays the distribution of calculated distances for the five hormone-bound complexes and apo receptor. The lower bounds for each box in **C** and **D** correspond to the apo receptors. **E)** Conformations obtained from accelerated MD simulations of unliganded AncSR2 were co-clustered with frames obtained from M75 mutants. M75A reveals substantial conformational overlap with AncSR2, M75F reveals no AncSR2 overlap while both M75L and M75I stabilize a minor conformational substate of AncSR2. **F)** For each M75 mutant, frames from progesterone- and estrogen-bound simulations were co-clustered with conformations from the unliganded simulation of the same receptor. In M75A, both hormones introduce new conformational states, suggesting that both would activate the receptor. M75F conformations are influenced by progesterone but not estradiol. M75I shows identical patterns between estradiol and progesterone, suggesting that the receptor would respond the same way to both hormones. M75L shows a similar pattern to M75I.

Next, we computed the H3-H5 interhelical distance in all variants by measuring Cα-Cα distances between two H3/H5 pairs: Q41/res75 and L42/W71 (**Fig. 2C, D)**. Importantly, while the Ala sidechain is not able to form van der Waals contacts with any H3 residues, all other position 75 substitutes were bulky enough for contact (Data not shown). M75A and M75I complexes show the smallest interhelical distances (7-7.2 Å), significantly shorter than other variants (**Fig. 2D)**. M75F shows the largest H3-H5 distance (7.5-8.1 Å) while M75L and WT AncSR2 show intermediate distances (7.4-8.1 Å). Thus, we determined that while M75 mutations preserve the contact with hormones, they vastly modulate the H3-H5 interhelical distance which might impact transcriptional activity.

### MD simulations predict that ligands selectively shift conformational states in AncSR2 variants

Our previous work indicated that NR conformational ensembles generated by MD simulations experience conformational shifts upon addition of ligand that reflect the activation potency of ligands. To characterize these ligand-induced effects in our engineered receptors, we used accelerated MD simulations to achieve enhanced sampling of the conformational space for each receptor-hormone combination. By lowering the energetic barrier for conformational transitions during simulations, this method allows us to visualize conformational states that may not be sampled in classical MD. For each variant, we obtained 500 ns accelerated MD trajectories for apo, estradiol- and progesterone-bound complexes.

To compare how the M75 mutations alter AncSR2 conformations, we performed combined clustering using frames (i.e., snapshots) obtained from the apo-AncSR2 trajectory with frames obtained from each of the apo-M75 mutants. Previously, we observed clear clustering differences between estrogen- and 3-ketosteroid-bound WT AncSR2 [38]. Estradiol-bound AncSR2 overlapped largely with apo-AncSR2, indicating that in the presence of an inactive ligand, AncSR2 samples the same conformations as it does in the apo state. Conversely, the addition of a 3-ketosteroid eliminated conformational overlap with the apo receptor, suggesting substantial ligand-induced conformational shifts [38]. The results show that mutations have strikingly distinct effects (**Fig. 2E)**. M75A shows substantial conformational overlap with WT AncSR2, as all clusters containing M75A snapshots also contained large numbers of WT frames. This result suggests that the M75A mutation does not have a huge impact on the conformational state of WT AncSR2. We observe the opposite trend when comparing M75F to WT: both complexes cluster into two unique (non-overlapping) clusters, suggesting that this mutation induces a large conformational effect in AncSR2. M75I and M75L reveal very similar patterns to one another: while the mutant receptors are largely retained in one cluster, the wildtype complexes segregate into 4-5 smaller clusters **(Fig 2E)**. This result suggests that a minor conformational state from WT AncSR2 is stabilized by these mutations.

Next, we used combined clustering to determine how progesterone and estradiol binding alters conformations within each variant (**Fig. 2F)**. We co-clustered frames from progesterone and estradiol-bound trajectories with frames from the corresponding apo simulation, allowing us to visualize differential conformational shifts induced by the binding of a 3-ketosteroid or an estrogen. In M75A, progesterone complexes form non-overlapping clusters with apo-M75A, while estradiol-M75A shows large apo overlap. Because progesterone shifts the conformational ensemble and estradiol does not, this observation would predict that the M75A variant is transcriptionally activated by progesterone but not estradiol. In contrast to M75A, estradiol-M75F fully overlaps with apo-M75F, while prog-M75F both induces new conformational states and retains small overlap with apo-M75F **(Fig. 2F)**. This result predicts that similar to WT AncSR2, estradiol would be unable to activate M75F while progesterone activates the mutant receptor weakly, compared to WT.

Interestingly, M75I shows nearly identical clustering patterns in both progesterone- and estradiol-bound complexes **(Fig. 2F**, M75I). In both cases, the ligand-bound receptor comprises one large cluster while the apo receptor fragments into five smaller clusters, suggesting that ligand binding is stabilizing a very minor conformational sub-state from the apo-M75I trajectory. Remarkably, M75L also shows very similar clustering patterns between its progesterone and estradiol-bound complexes **(Fig. 2F**, M75L). In both cases, the ligand-bound trajectory fragments into two large clusters, as does the apo trajectory. Overall, both M75I and M75L variants show ligand-induced conformational shifts, predicting activation by both hormones for both receptors. Because the response is nearly identical in both hormones, another possible interpretation of these clustering patterns is that the receptor will respond the same way to both ligands, which may also suggest that the receptor has ligand-independent function, i.e. either inactive or constitutively active.

### Transcriptional responses in M75 variants span a broad activity spectrum

To compare the predicted conformational shifts from MD studies with transcriptional function in AncSR2 variants, we measured transactivation in cell-based luciferase reporter assays using the five aforementioned hormones. All 3-ketosteroid hormones activate AncSR2 with EC_50_ in the sub-nanomolar range except DHT which had a nanomolar EC_50_, consistent with earlier reports [39] (**Fig. 3A)**. As previously observed, estradiol is not able to activate the receptor (Table 1 (**Fig. 3G**)).

**Figure 3:**
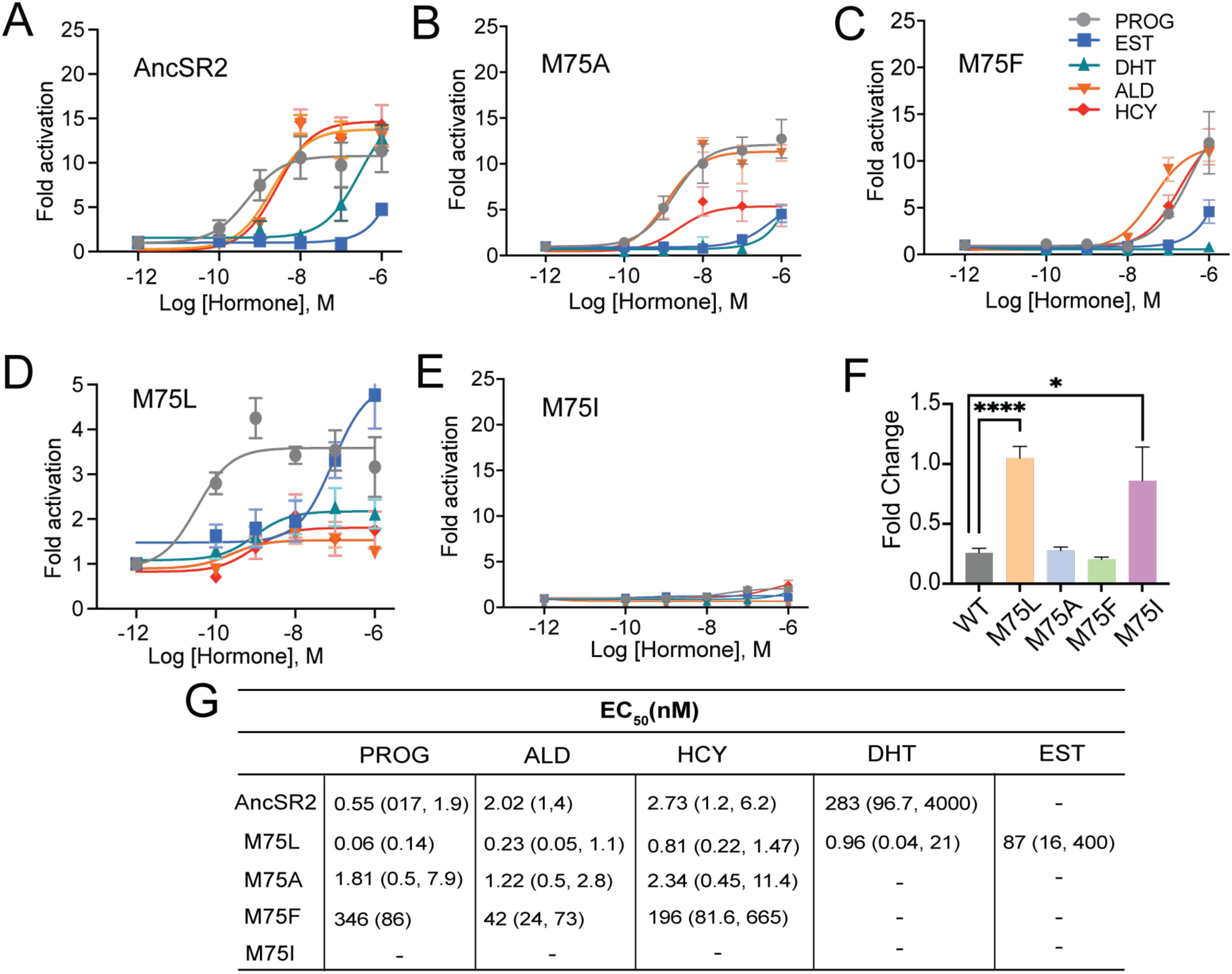
Differential ligand activation of AncSR2 variants. Dose-response curves of AncSR2 and its variants in the presence of aromatized and non-aromatized hormones. **A)** AncSR2 receptor showed differential response to non-aromatized hormones with no response to estradiol. **B)** M75A substitution slightly increased the fold activation as compared to AncSR2 with no change in the receptor efficacy for progesterone, aldosterone, hydrocortisone. **C)** M75F substitution decreased the receptor responsiveness for hormones. **D)** M75L receptor efficacy is significantly reduced compared to WT AncSR2. In contrast to WT AncSR2, M75L was activated by estradiol. **E)** M75I substitution completely abolished the ligand activation for both types of hormones. Each data point is an average of two to three biological replicates. The error bar associated with each data point represent SEM. **F)** M75L and M75I receptors fold change over empty vector in the absence of hormone suggest that they exhibit constitutive activity. Two-tailed unpaired t-test, (****) P < 0.0001, (*) P < 0.05. **G)** Table shows EC_50_ values obtained from the analysis of dose-response curves of individual receptors for different hormones. 95 % confidence interval values are shown in parentheses.

In the M75 mutants, we observe a wide range of functional behavior that is remarkably consistent with predictions from clustering of MD-generated ensembles. The M75A variant largely recapitulates the activity profile of WT with the largest differences being loss of DHT activation and reduced efficacy (E_max_) in aldosterone (**Fig. 3B)**. Importantly, M75A is activated by progesterone and unresponsive to estradiol, as predicted in simulations. Potency is reduced for all ligands in M75F activation **(Fig. 3C)**, with no activity observed in estradiol and DHT. Progesterone activation in M75A is reduced by ∼3 orders of magnitude relative to WT AncSR2, consistent with the prediction of weaker activation based on partial conformational shift observed in simulations.

None of the hormones activated M75I **(Fig. 3E)**, suggesting that this mutation may inhibit ligand binding. Strikingly, while the M75L variant is the only receptor activated by aromatized and non-aromatized hormones, the efficacies of the hormones are drastically reduced (E_max_ ∼ 2-5) **(Fig 3D)** compared to AncSR2 (E_max_ = 11-16) **(Fig. 3B)**, suggestive of reduced response in M75L. We then assayed transactivation in the absence of ligands for all AncSR2 variants and confirmed the presence of basal activity in the M75L mutant **(Fig. 3F)** that is independent of the cell lines used **(Fig. S2)**. Thus, both M75I and M75L variants introduce a new interpretation of the clustering results. The observation of a nearly identical response to both hormones indicates that the receptor is agnostic to the identity of the ligand and will produce the same functional response to either.

### Ligand binding and overall AncSR2 dynamics are impacted by M75 mutations

M75I and M75L mutants displayed unexpected results in transcriptional assays. To understand the molecular basis for these effects, we developed a binding assay to probe hormone binding to AncSR2, M75L and M75I variants. For this purpose, we have designed a probe by linking 11-deoxycorticosterone (11-DOC), a potent AncSR2 agonist [39] to fluorescein (FAM) (See Methods). By titrating purified AncSR2 LBD against a fixed concentration of 11-DOC-FAM **(Fig. 4A)**, we obtained a saturation binding curve with an equilibrium dissociation constant K_d_ = 180 nM **(Fig. 4B)**. To validate that 11-DOC-FAM binds the AncSR2 ligand binding pocket, we used a competition assay to measure the K_i_ (inhibition constant) of unlabeled 11-DOC. We observe that unlabeled 11-DOC outcompeted the 11-DOC-FAM with K_i_ = 33 nM, approximately five-fold lower than K_d_ (**Fig. 4C**). Similarly, previous fluorescent probes for SRs have been reported with 10-fold lower K_d_ compared to the unlabeled ligand [42].

**Figure 4:**
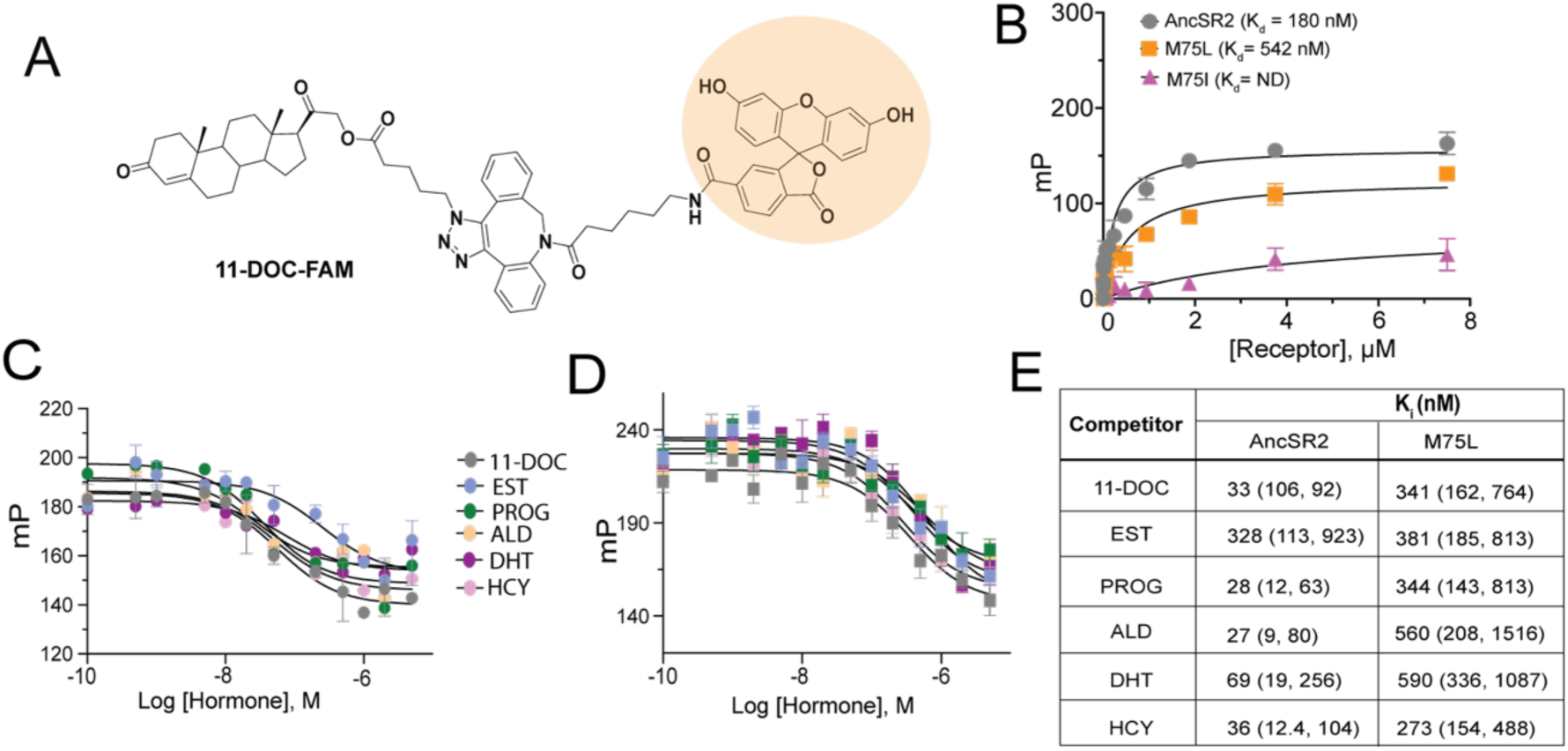
Fluorescence polarization assay of binding affinity of AncSR2 and M75 mutants. **A)** Structure of synthesized probe: FAM labelled 11-DOC (11-DOC-FAM). **B)** 11-DOC-FAM binds to AncSR2-LBD and M75L with equilibrium dissociation constant, K_d_=180 nM (70, 425) and K_d_= 542 nM (237,1155), respectively. 95% confidence intervals shown in parentheses. Binding data is the average of six (for WT AncSR2 and M75L) and three (M75I) replicates from two and one independent experiment, respectively where each experiment consists of three independent replicates. **D)** and **E)** FP-based competition binding experiments shows that all steroid hormones bind WT AncSR2 and M75L with nM inhibition constants (K_i_). Five hormones used in this study: progesterone (PROG), aldosterone (ALD), hydrocortisone (HCY), 11-deoxycorticosterone (11-DOC), Dihydrotestosterone (DHT) and Estradiol (EST) (chemical structures shown in **Fig. S3**). Error bars indicate SD from three independent replicates. E) K_i_ values obtained for five hormones from FP-based competition ligand binding assay. 95% confidence intervals from independent triplicate measurements are shown in parentheses.

We then used the competition assay to determine K_i_s for four 3-ketosteroids (progesterone, DHT, hydrocortisone, aldosterone) and estradiol (**Fig. 4C**). With varying affinities, all ligands are able to outcompete 11-DOC-FAM from the AncSR2 binding pocket (**Fig. 4E)**. The K_i_ values for progesterone, hydrocortisone and aldosterone only differed slightly, but were lower than those observed for DHT and the aromatized hormone, estradiol **(Fig. 4E**). Thus, AncSR2 binds 3-ketosteroids preferentially over aromatized hormones, with our results suggesting that a C17 acetyl substituent in hormones may confer a binding advantage. Using the same assay for M75L, we observe that compared to WT AncSR2, the M75L substitution reduces the receptor’s affinity for 11-DOC-FAM **(Fig. 4B)** and all steroid hormones (**Fig. 4D)**. None of the ligands bind M75I, which explains why this mutant was not activated by the steroids **(Fig. 4B, Fig. 3E)**.

To learn how the M75L variant is constitutively active, we performed HDX-MS to probe structural and dynamical changes in the mutant at fast deuterium exchange time scales (t=1-60 min). With peptide coverage ranging from 85-89% **(Fig. 5A)** for the AncSR2 and M75L LBDs, we monitored the dynamics of nearly the entire protein at different time points **(Fig. S4-S7)**. First, we identified the effects of the M75L mutation on WT AncSR2 dynamics using a comparative HDX (ΔHDX) analysis **(Fig. 5B)**. Higher deuterium uptake was observed in multiple regions in the M75L mutant, indicating that the mutation broadly destabilized the LBD. Deprotection is observed in residues surrounding the ligand binding pocket, including H3, H7, H10, H12, but unexpectedly in distant regions such as H9.

**Figure 5:**
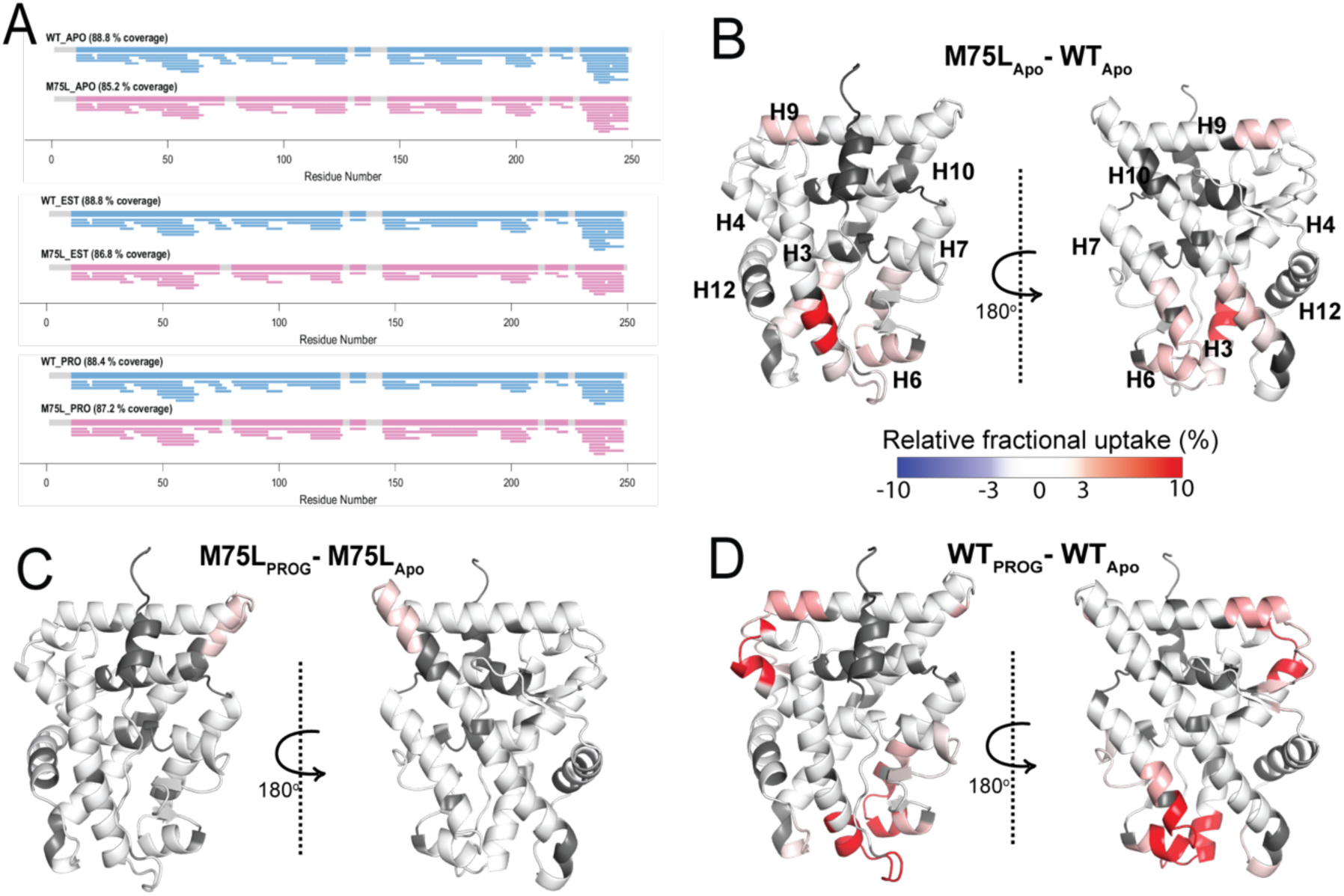
Conformational effects of AncSR2 M75L mutation probed by HDX-MS. **A)** Sequence coverage maps for WT AncSR2 (blue) and M75L (pink) in their apo, estradiol (EST), and progesterone (PRO) bound form. **B-D)** The percent difference in relative fractional uptake (ΔRFU) of deuterium between **B)** M75L and WT AncSR2 apo states (M75L_Apo_ – WT_Apo_), **C)** M75L-progesterone bound and M75L Apo states (M75L_PROG_-M75L_Apo_) and **D)** WT-progesterone bound and WT Apo states (WT_PROG_-WT_Apo_). All plots represent a 15 min time point. Color bar indicates the fractional difference in relative deuterium uptake for the two states compared. Positive numbers (red) indicate higher deuterium exchange in state A relative to state B (i.e. deprotection) while negative numbers (blue) indicate lower deuterium exchange (i.e. protection). Dark grey regions represent peptides with no sequence coverage.

To explore the dynamic effects of the M75L mutation on ligand binding, we analyzed ΔHDX comparing the progesterone-bound forms to their apo counterparts for both WT and M75L receptors **(Fig. S5)**. While very little change in deuterium exchange accompanied progesterone binding in M75L **(Fig. 5C)**, dramatic enhancement in dynamics was observed across AncSR2, including peptides ^20^AGYDNTQPDTTNYLL^34, 48^VVKWAKALPGFRNLHLDD^65, 104^NEQRMQQSAM^113, 145^LLSTVPKEGLKSQ^160^, suggesting that the two variants are differentially affected by the addition of progesterone **(Fig. 5D)**. A similar effect was seen in the estradiol-bound forms of the receptors (**Fig. S5)**. WT AncSR2 showed deprotection in several regions when bound to estradiol, while M75L showed no changes in deprotection/protection. These experimental observations strongly support a model in which the M75L mutation shifts the ensemble conformation to a ligand-bound state, allowing the receptor to be less dynamically responsive to the addition of ligand. This effect may also explain the reduced ligand binding ability observed in the M75L variant.

### Conformational analysis of MD trajectories

To describe the molecular mechanisms underlying the conformational shifts observed in MD trajectories, as well as understand how the M75 mutations achieve the range of functional profiles observed in luciferase and binding assays, we sought to visualize the conformational effects that accompany M75A, M75F and M75I substitutions. By performing a close examination of our MD trajectories of the M75A variant, we observed two new interactions that potentially stabilize hormones, formed by Trp71 (H5) and Gln41 (H3) **(Fig. 6A)**. Both residues are conserved in SRs: W71 mediates interactions between bound ligands and H12 [43-45] while Q41 is positioned to stabilize the A-ring of hormones. Simulations predict that reduced bulk at position 75 allows both W71 and Q41 sidechains to gain proximity to the hormone and potentially provide increased stability. In support of this hypothesis, a Q41A mutation in the background of M75A abolished activation by 3-ketosteroids **(Fig. S8)**.

**Figure 6.**
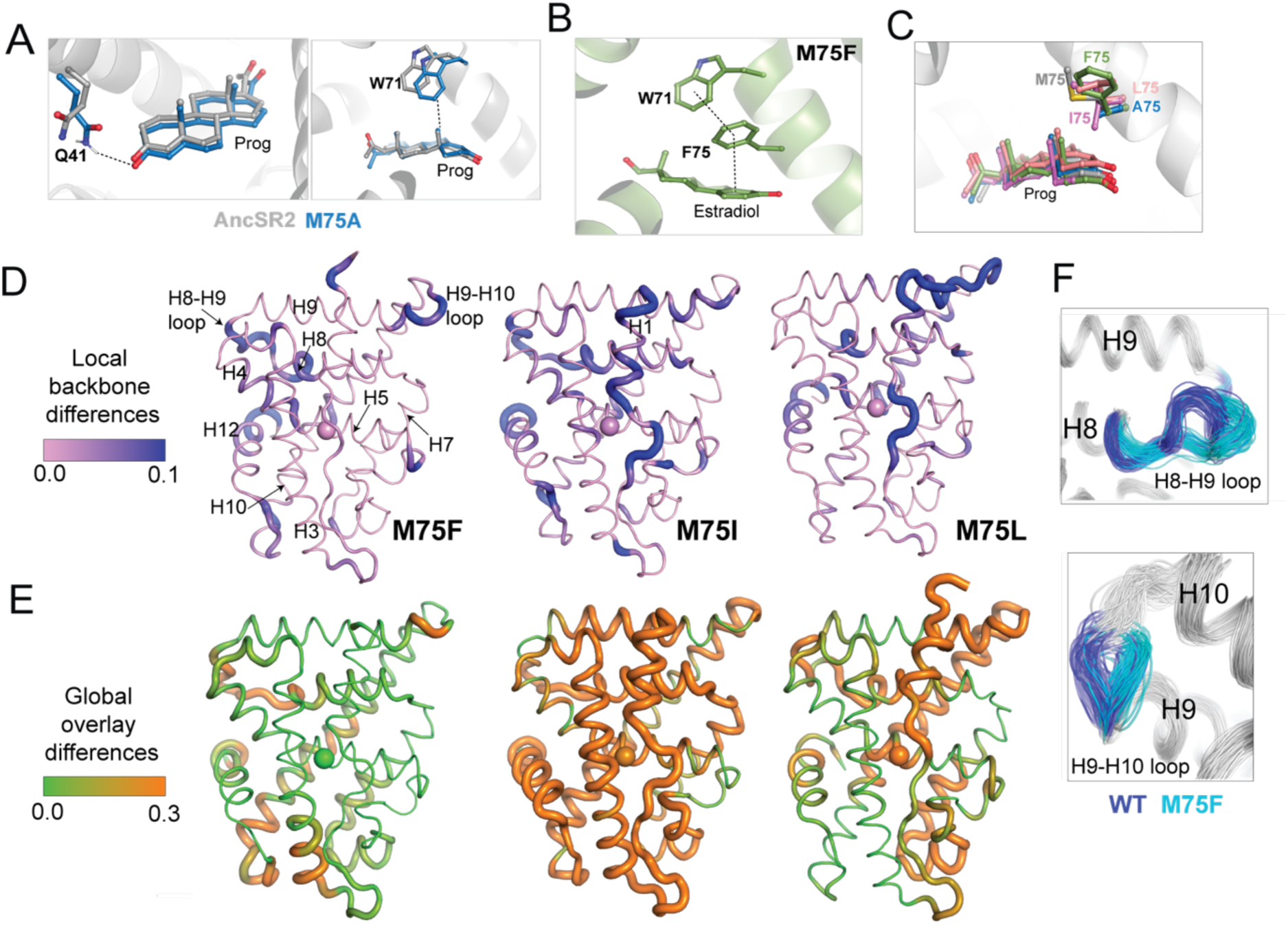
Structural analysis of MD simulations of AncSR2 variants to explain altered transcriptional response. Two potential explanations for enhanced activity in **A)** M75A are the proximity of Q41 (top) and W71 (bottom) to hormones, due to the drastically reduced H3-H5 distances. **B)** In estradiol-bound M75F, a pi-stacking triad is formed between W71, F75 and the hormone A-ring. This observation suggests an explanation for why the mutant is selectively activated by 3-ketosteroids but not by estrogens. **C)** Simulations predict that the Ile sidechain of M75I is likely to insert into the binding pocket and interfere with ligand binding. This effect is not observed any of the other complexes. **D)** Ensemblator analysis of local/backbone changes of M75 variants compared to WT AncSR2. Conformational changes are quantified and colored by the discrimination index (DI). Lower DI values indicate regions where structures are nearly identical between WT and mutant receptors while higher DI indicates changes in local backbone angles induced by mutations. Spheres on H5 identify position 75. **E)** Ensemblator analysis of global changes in M75 mutants compared to WT. Structures are globally overlaid prior to analysis. Structures are colored by DI, where lower values correspond to highly similar regions in superimposed structures while higher values indicate regions of structural dissimilarity. Spheres on H5 identify position 75. **F)** Overlay of 200 structures used for Ensemblator analysis showing conformational changes observed in H8-H9 and H9-H10 loops. Structures from WT AncSR2 are colored blue while mutant structures are shown in cyan, illustrating that structural variations are the result of mutation.

In M75F, we observe from simulations that while F75 preserves a van der Waals contact between the two helices, the bulkiness of the sidechain gives M75F complexes the largest interhelical distances **(Fig. 2B, C)**. Additionally, F75 engages W71 in a hydrophobic interaction. We measured the fraction of time that the aromatic sidechains engage in pi-stacking and interestingly, the occupancy was less than 5% for 3-ketosteroid complexes but rose to ∼40% in the M75F-estradiol complex. Additionally, W71, F75 and the aromatic A-ring of estradiol form a triad in this complex **(Fig. 6B, C)**, which was absent in 3-ketosteroid complexes. Thus M75F bears strong similarities with AncSR2, where the Phe sidechain, similar to the WT Met, engages in pi interactions with estrogens but not 3-ketosteroids [38]. Unsurprisingly, while potency is reduced by 1-2 orders of magnitude, M75F displays a similar activation profile to WT AncSR2 (**Fig. 4A, C**). Notably, DHT does not activate M75F in our assay, which we believe results from the same reason that DHT weakly activates AncSR2, i.e., the lack of a C17 acetyl substituent to stabilize the D-ring end of the hormone via hydrogen bonding. Overall, results suggest that the increased H3-H5 distance resulting from the bulky Phe substitution is responsible for the suboptimal functional profile of this mutant. In the M75I variant, consistent with the lack of ligand binding and transcriptional activation observed, simulations show that the beta-branched Ile sidechain is positioned to enter the binding pocket and interfere with ligand binding (**Fig. 6D, 4C**).

Finally, we sought to reveal the conformational features defining the ensembles generated from M75 mutants. We focused on unliganded M75F, M75I and M75L, as these three variants revealed a conformational shift when clustered with WT AncSR2 (**Fig. 2E**). To achieve this conformational analysis, we obtained 100 representative structures each from the largest WT and mutant clusters of **Fig. 2E**. We compared the subgroups using Ensemblator [46, 47] to quantify both differences and similarities between mutant and WT populations. This method achieves local conformational comparison by calculating the similarity between backbone conformations based on dihedral angles [47]. Global comparisons are performed based on an atom-level overlay of all 200 structures. Comparisons are represented by Discrimination Index (DI), a metric ranging between 0 and 1 that reveals the most significant local and global differences between both subgroups (See Methods).

Conformational changes are colored by calculated DI, where higher values identify structural differences in the mutant relative to WT (**Fig. 6D, E**). Here, we observe small changes in local (< 0.3) and global DI (< 0.5) for most regions of the receptors (**Fig. S9**), indicating that subtle conformational changes drive the structural effects detected via clustering. Thus, we have focused on analyzing these subtle differences between mutant and WT AncR2 structures. In all variants, M75 mutation induces a backbone change in H5 beginning at M72 (M75I) or W71 (M75F, M75L). These changes propagate to the adjacent H8, subsequently influencing the H8-H9 loop and/or the H9-H10 loop (**Fig. 6F**). Other regions impacted include H10 and the pre-H12 loop. Global changes vary more drastically between the three variants. M75F shows the largest effects in H8, H10 and the bottom of H3. Conversely, M75I undergoes global shifts > 0.3 DI in nearly all helices, while M75L is most affected at H7, H8 and H10. Strikingly, we note that the largest global changes in M75L coincide with the regions predicted by HDX-MS to be destabilized by the M75L mutation (**Fig. 5B**).

## Discussion

Here, we have used molecular dynamics simulations to generate conformational ensembles of NR complexes, with a goal of revealing how ligand-induced population shifts correlate with the functional properties of ligands. We engineered AncSR2 variants with altered potency and specificity by substituting M75, a known regulator of activation and ligand recognition in steroid receptors. Our results reveal a strong relationship between ligand potency and the extent of population shifts in ligand-bound ensembles relative to apo NR ensembles. Clustering shows that active ligand complexes do not overlap with unliganded complexes while inactive ligands result in complete overlap, suggesting that the addition of ligand does not shift the receptor out of an inactive, ligand-free state. Notably, because the M75 mutations do not perturb the AF-2 surface, the ensembles reflect extremely subtle conformational changes originating at H5 and thus highlight the exquisite sensitivity of this computational assay.

Unexpectedly, we also observe partial overlap in clustering for the M75F mutant, which we attribute to a partial switch of the conformational ensemble, associated with partial agonism or reduced activation potency. Future studies will be important to determine whether MD-generated ensembles of NR ligand complexes can be closely parsed to distinguish between partial agonist (low-efficacy) vs weak agonists (low-potency). Fortuitously, the M75 mutants selected in our work allowed us to access a constitutively active receptor as well as an inactive variant. Conformational ensembles generated for these complexes were particularly intriguing, as they predicted that aromatized and non-aromatized hormones would induce identical population shifts in the receptor, suggesting that the receptors were agnostic to the functional profile of the ligand.

By combining a structural analysis of our MD trajectories with biophysical experiments, we determined the mechanisms underlying the altered functional profile of each NR variant. While we anticipated a loss-of function with the M75A mutant due to the elimination of interhelical H3-H5 contacts, we observed that the variant retains activity similar to the WT receptor. An explanation that emerges from our MD simulations is that one or both of two conserved pocket residues, Trp71 and Gln41 may gain proximity to the hormone and provide stabilization. Another contributing factor, based on previous SR studies is that the lack of a bulky sidechain at position 75 may create excess volume in the binding pocket, allowing ligands to bind without perturbing H12 or the AF-2. Such a state would explain the constitutive activity and/or gain of function observed in M75A mutants of PR and MR [41].

Conversely, the larger Phe substitution introduces the largest H3-H5 distances in our simulations (**Fig. 2B, C**). As there are no other structural changes observed across the variant, we attribute the reduced potency for all hormones and the lower thermodynamic stability in M75F to the increased interhelical distance. However, simulations reveal that the Phe sidechain distinguishes between aromatized and non-aromatized hormones via pi interactions, confirming the role of position 75 in mediating ligand specificity. These pi interactions with estradiol cause frustration around the A-ring, preventing stabilization and activation of WT AncSR2 and M75F by estrogens. Additionally, our results support the previously established importance of steroid D-ring interactions with the conserved H3 residue, N37 [48-50]. Because neither estradiol nor DHT activate M75F and both lack the ability to interact with N37, this residue could be playing a compensatory role in this impaired variant, permitting activation by aldosterone, hydrocortisone and progesterone.

The most unexpected result was the constitutive activity observed in the M75L mutant, surprising because the variant retained similar stability as the WT receptor with only slight differences in secondary structure (**Fig. 1C,D**). Ligand binding assays showed that M75L has reduced affinity for our 11-DOC-FAM probe compared to WT AncSR2, as well as a lower measured K_i_. Increased conformational dynamics around the ligand binding pocket in the apo M75L receptor compared to WT in HDX-MS studies suggests that the mutation likely affects ligand binding, supported by reduced K_i_s observed in binding assays.

However, the absence of a change in ΔHDX upon addition of ligand to M75L supports our claim that the receptor adopts a partially active state that mimics ligand-bound AncSR2, resulting in low levels of constitutive activity and a reduced response to ligands.

This work presents a highly sensitive *in silico* approach for describing ligand-specific conformational changes in NR ensembles. Because of the demonstrated importance of NR ensembles for understanding and predicting activity profiles of ligands, our findings confirm the inherent promise in the use of MD-generated ensembles as a predictive tool in ligand design. An added advantage is that analysis of MD trajectories is useful for providing a structural description of the ensemble, as well as elucidating the mechanisms by which ligands induce distinct conformational effects in NRs. We have used Ensemblator to perform reveal the local and global structural perturbations associated with three of the mutations investigated. We also observe that M75 mutations induce dynamic effects at distant regions of the receptor, including helices 9 and 10, which will be explored in future work to determine the potential for functional modulation of receptors via novel mechanisms.

## Materials and Methods

### Materials

Sodium chloride, Trizma, glycine, sodium dodecyl sulphate, ethylenediaminetetraacetic acid (EDTA), imidazole, glycerol were purchased from Fischer scientific (USA). Ampicillin, tryptone, yeast extract, isopropyl β-D-1 thiogalactopyranoside (IPTG) were procured from RPI chemicals (USA). DTT was purchased from Alfa Aesar, USA. Estradiol and ammonium persulphate were purchased from Sigma Chemical Co. (USA). Hydrocortisone, 11-Deoxycorticosterone (11-DOC), progesterone, Dihydrotestosterone, 11-Deoxcycortisterone acetate were purchased from Acros organics. Bioscale Nuvia-IMAC Ni-charged and ENrich SEC70 and SEC650 10/300 size exclusion columns were purchased from Biorad, USA and used with BioRad NGC Quest plus FPLC system. Syringe filters (0.2 micron) were procured from Millipore corporation. All reagents and chemicals were of analytical grade

### Cloning, expression, and purification of ligand binding domain of WT and mutants

The gene encoding AncSR2 was PCR amplified from the vector pSG5-Gal4-DBD-SR2-LBD using forward and reverse primers containing *Eco*RI and *Hind*III restriction sites, respectively, to clone into pMALCH10T vector. Primers were designed (**Table S1**), followed by mutagenesis to generate M75L, M75A, M75I and M75F mutants of AncSR2-LBD in both pSG5 and pMALCH10T vectors. Mutants were confirmed by DNA sequencing.

MBP-His-tagged LBDs of WT-AncSR2 and its mutants were expressed in and purified from *E. coli*. BL21(DE3) as previously reported with slight modification [39]. Briefly, cells containing the respective plasmids were grown in LB broth till O.D_600_ reaches 0.6-0.8 at 37 °C. The protein expression was induced by addition of 0.3 mM IPTG and 50 μM progesterone and grown further at 30 °C for 4 hours. The cells were harvested by centrifugation at 8000 RPM for 10 minutes. The cells were lysed by sonication using a 10 sec pulse-on and 30 sec pulse-off cycle. Cell debris was removed by centrifugation of lysate at 15000 RPM for 40 minutes. The cleared supernatant was purified by Ni-Affinity chromatography (Nuvia-IMAC). The purified protein was subjected to TEV protease treatment (0.5 mg/ml) overnight and simultaneously dialyzed against buffer containing 20 mM Tris (pH 7.4), 150 mM NaCl and 10 % glycerol. The dialyzed lysate was twice purified by a Ni-affinity column, followed by collection of flow through containing desired LBDs. LBDs were finally purified by SEC70 gel filtration column using Bio-Rad NGC plus system in a buffer containing 20 mM Tris (pH 7.4), 150 mM NaCl and 10 % glycerol and stored in small aliquots at -20 °C until further use. It should be noted that all steps from cell harvesting to gel filtration were either performed on ice or at 4 °C, unless stated otherwise. The purity of the purified protein was assessed by loading the sample on the 14 % SDS-PAGE.

For ligand binding assays, MBP-tagged AncSR2 LBD and mutants were expressed and purified (without protease treatment) as described above for the LBD, except for the use of 0.4 mM IPTG and 50 μM 11-DOC acetate for induction of protein expression, followed by overnight growth at 18 °C. The MBP tagged protein was purified by Ni-affinity chromatography, followed by gel filtration in pH 7.4 buffer containing 20 mM Tris (pH 7.4),150 mM NaCl and 10 % glycerol. The final purification was performed using a SEC650 gel filtration column in a buffer containing 20 mM HEPES (pH 7.4), 150 mM NaCl, 3 mM EDTA, 5 mM DTT and 0.005 % Triton X-100.

### Circular dichroism measurements

Far UV-CD measurements were done on a Jasco J-1500 spectrophotometer equipped with a temperature controller. The far-UV CD spectra of the WT and its mutants were measured in the wavelength range 195-250 nm. For spectral measurements,1 mm path length cuvette was used, with scan rate of 50 nm/sec, 1 sec response time and bandwidth of 1 nm and protein concentration used was 0.2 mg/ml. The CD instrument was continuously purged with N_2_ gas at 5-8 lit/min flow rate and routinely calibrated with D-10-camphorsulfonic acid. Each spectrum was an average of 3 consecutive scans and corrected by subtraction of the buffer (10 mM phosphate, pH 7.4 and 100 mM NaCl). The raw CD data was converted into mean residue ellipticity at a wavelength, [*θ*]_λ_ (deg cm^2^dmol^-1^) by using the relation,

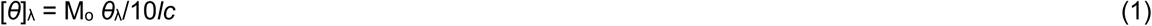

where, *M*_o_ is mean residue weight of the protein, *θ*_λ_ is the observed ellipticity in millidegrees at λ wavelength, *c* is the concentration of protein in mg ml^-1^, and *l* represents the cuvette path length in centimeters.

### Thermal denaturation measurements

Thermal denaturation was followed by measuring changes in the CD signal at 222 nm as a function of temperature. The heating rate of 1°C with bandwidth 4 nm, 2 sec response time was used in the temperature range 20-70 °C to follow the denaturation. The raw CD data was converted into the concentration independent parameter ([*θ*]_λ_) using equation 1. In the analysis of the denaturation curves,a two-state model (N = native state, D = denatured state) was assumed, and the temperature dependencies of pre- and post-denaturation baselines are linear. Stability curves (Δ*G* vs temperature) were constructed choosing values of Δ*G* (± 1.3 kcal/mol) close to the midpoint of denaturation (*T*_m_) that fall on the straight line. A linear least square analysis was used to estimate the entropy change at *T*_m_ (= - δΔ*G*/δT)_p_ which is then multiplied by *T*_m_ to get the values of apparent Δ*H*_m_. It should be noted that the heat-induced denaturation process was irreversible in the measured experimental conditions, so the stability parameter is defined here as apparent *T*_m_.

The fraction of denatured molecules (*f*_D_) was calculated by the relation:

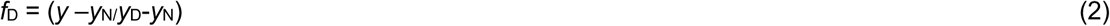

where y is the observed optical property of protein at temperature *T, y*_N_ and *y*_D_ are optical properties of native and denatured molecules at the same temperature.

### Ligand binding and competition assay

WT-AncSR2 and the M75L mutant were expressed and purified as MBP-tagged proteins. All fluorescence polarization experiments were performed buffer containing 20 mM HEPES (pH 7.4), 150 mM NaCl, 3 mM EDTA, 5 mM DTT and 0.005 % Triton X-100. For saturation binding experiments, the binding affinity (K_d_: dissociation constant) of the receptor for the probe dexamethasone-fluorescein (11-DOC-FAM) was determined using a constant concentration of 10 nM of the probe and a variable receptor protein concentration of 7.5 x10^−6^ – 4.5 x10^−10^ M in a 384 well plate. The plate was centrifuged at 500 RPM for 2 minutes and incubated overnight at 4 °C before reading. Fluorescence polarization measurements were performed on a Spectramax iD5 plate reader (Molecular Devices, USA) using excitation and emission wavelengths of 485 and 528, respectively. Six technical replicates and two biological replicates were obtained and data was plotted as the average mP (millipolarization) of all replicates versus receptor protein concentration. The saturation binding curve was analyzed by a one-site hyperbola binding model using GraphPad Prism vs 9 (GraphPad, Inc, La Jolla, USA). In the competition binding assay, 10 nM 11-DOC-FAM and protein concentration approximately 1.1-1.8 times K_d_ for 11-DOC-FAM were incubated with variable (10^−10^–10^−5^ M(competitive ligand overnight at 4 °C. The observed mP values in the presence of competitive ligands were plotted and fit by the *Fit Ki* model of GraphPad Prism vs 9. All data points were plotted after buffer subtraction.

### Luciferase reporter assays

Hela cells were grown and maintained in phenol red free medium MEM-α supplemented with 10 % charcoal-dextran stripped FBS. Cells were seeded in 96-well plates at 70-90 % confluency and co-transfected with 1 ng Renilla (pRL-SV40), 50 ng 9x-UAS firefly luciferase reporter and 5 ng pSG5-Gal4DBD-LBD fusions of WT-AncSR2, M75L, M75A, M75I, and M75F receptor plasmids using FuGene HD (Promega). Cells were treated with DMSO or varying drug concentrations 24 hours after transfection, all in triplicate. Firefly and Renilla luciferase activities were measured 24 hours after drug treatment using Dual-Glo kit (Promega) using a Spectramax iD5 plate reader. Fold activation is represented as normalized luciferase over DMSO control. Dose response curves were generated by GraphPad Prism v9.0.

### Molecular dynamics simulations

Coordinates for AncSR2 ligand binding domain were obtained from PDB 4FN9, AncSR2 LBD-progesterone[39]. Ligand complexes with the other hormones in this study (dihydrotestosterone, hydrocortisone, aldosterone, estradiol) were constructed by manually modifying the steroidal core of progesterone. All waters and surface-bound molecules from the crystallization buffer were deleted from the models. M75 mutant AncSR2 complexes were prepared by using Xleap in AmberTools20 [51] to replace the M75 sidechain with alanine, leucine, isoleucine and phenylalanine respectively. In total, 30 complexes (5 AncSR2 variants, 5 hormones and 1 unliganded state per variant) were prepared for simulation.

All complexes were prepared using Xleap. Parameters for steroid hormones were obtained using the Antechamber [52] and the Generalized Amber ForceField [53]. The parm99-bsc0 [54]forcefield was used for protein residues. Briefly, complexes were solvated in an octahedral box of TIP3P water [55], allowing a 10 Å buffer around the protein. Na+ and Cl-ions were added to achieve a final concentration of 150 mM. Minimization was performed in four steps. First, 500 kcal/mol.Å^2^ restraints were placed on all solute atoms, and 5000 steps of steepest descent performed, followed by 5000 steps of conjugate gradient minimization. In the second step, this protocol was repeated with restraints reduced to 100 kcal/mol.Å^2^. Restraints were then removed from protein atoms and retained on the hormone for a third minimization step, followed by a final unrestrained minimization for all atoms.

Complexes were heated from 0 to 300 K using a 100 ps run with constant volume periodic boundaries and 5 kcal/mol Å^2^ restraints on all solute atoms. All simulations were performed using AMBER 2020 on GPUs[53, 56] Before production MD simulations, 10-ns simulations with 10 kcal/mol.Å^2^ restraints on all solute atoms were obtained in the NPT ensemble. This was followed by a second 10-ns simulation with restraints reduced to 1 kcal/mol.Å^2^. Finally, restraints were retained only on the ligand atoms for a third 10-ns equilibration step. Production trajectories were obtained on unrestrained complexes, each complex simulated for 500 ns in triplicate. All MD was performed with a 2 fs timestep and using the SHAKE algorithm [57] to fix heavy atom hydrogen bonds. Simulations were performed with the NPT ensemble, a cutoff of 10 Å to evaluate long-range electrostatics and particle mesh Ewald and van der Waals forces.

### Accelerated MD

Accelerated MD was used as previously reported [38, 58] to enhance conformational sampling for AncSR2 complexes. We apply a dual-boosting approach, selecting parameters for potential energy threshold (EP), dihedral energy threshold (ED), dihedral energy boost (aD) and total potential energy boost (aP) using published guidelines [59]. 500 ns accelerated MD simulations were performed for a total of 15 complexes (5 AncSR2 variants, 2 hormones and 1 unliganded state per variant). All simulations were performed in AMBER 2020 and classical MD simulations were used to obtain Average dihedral energy (EavgD) and average total potential energy (EavgP)

aD: 0.2 * (EavgD + 3.5 (kcal/mol *Nsr)

ED: EavgD + 3.5 kcal/mol * Nsr

aP: 0.16 kcal/mol * Natom

EP: EavgP + (0.16 kcal/mol*Natom) (where Nsr = number of total solute residues, Natom = total number of atoms)

#### Analysis

Structural averaging and analysis were performed with the CPPTRAJ module of AmberTools17 [60]. The ‘strip’ and ‘trajout’ commands of CPPTRAJ were used to remove solvent atoms and obtain twenty-five thousand evenly spaced frames from each simulation for analysis. For each complex, triplicate runs were combined to yield seventy-five thousand frames for analysis. Two residues were defined to be within in Van der Waals contact if the distance between a pair of heavy atoms from each residue was < 4.5 Å. The ‘distance’ command of CPPTRAJ was used to obtain these distances.

#### Clustering and Conformational Analysis

The MMTSB toolset [61] was used to perform clustering of accelerated MD trajectories. clustering analyses. For each complex, 25,000 evenly spaced conformations were obtained from each 500 ns trajectory for clustering. A 2.4 Å cutoff was used. Clustering was performed between mutant and WT variants, as well as between liganded and apo complexes.

### Amide Hydrogen Deuterium Exchange Mass Spectrometry

Wild type, M75L, and M75I SR2 samples were stored in 20 mM Tris, 150 mM NaCl, pH 7.4. To assess allostery in response to ligand binding (estradiol and progesterone), 10 μM SR2 samples were incubated with 200 µM ligand at 37 °C for 150 minutes before HDX. Deuterium labelling was carried out using a PAL RTC autosampler (LEAP technologies). All samples were diluted to a final concentration of 90.9% D_2_O to initiate the deuterium exchange reaction. Deuterium buffers were prepared by dilution of 20X storage buffer in D_2_O. Deuterium exchange was carried out at room temperature (20 °C) maintained on a drybath for 10, 30, 60, 900, and 3600 sec followed by rapidly quenching the reaction to minimize back exchange using 1.5 M GdnHCl and 0.1% FA on ice to bring the pH down to 2.5.

Quenched samples were injected onto an immobilized pepsin treatment (BEH Pepsin Column, Enzymate, Waters, Milford, MA) using a nano-UPLC sample manager at a constant flow rate of 75 µl/min of 0.1% formic acid. Proteolyzed peptides were then trapped in a VanGuard column (ACQUITY BEH C18 VanGuard Pre-column, 1.7 µm, Waters, Milford, MA) and separated using a reversed phase liquid chromatography column (ACQUITY UPLC BEH C18 Column, 1.0 × 100 mm, 1.7 µm, Waters, Milford MA). NanoACQUITY binary solvent manager (Waters, Milford, MA) was used to pump an 8-40% acetonitrile gradient at pH 2.5 with 0.1% formic acid at a flow rate of 40 µl/min and analyzed on a SYNAPT XS mass spectrometer (Waters, Milford, MA) acquired in MS^E^ mode [62].

Undeuterated SR2 particles were sequenced by MS^E^ to identify pepsin digested peptides using Protein Lynx Global Server Software (PLGS v3.0) (Waters, Milford, MA). The peptides were identified by searching against the SR2 protein sequence database with a non-specific proteolysis enzyme selected. Peptides from the undeuterated samples that were identified and matched from the primary sequence database were filtered and considered with the following specifications: precursor ion tolerance of < 10 ppm, products per amino acid of at least 0.2 and a minimum intensity of 1000.

Average deuterium exchange in each peptide was measured relative to undeuterated control peptides using DynamX v3.0 (Waters, Milford, MA) by determining the centroid mass of each isotopic envelope. Subtractions of these centroids for each peptide from the undeuterated centroid determined the average number of deuterons exchanged in each peptide [63]. Deuterium exchange for all peptides is represented using relative fractional uptake (RFU) plots. Each value reported is an average of three independent deuterium exchange experiments and not corrected for back-exchange [62]. Difference plots were made by subtracting absolute centroid mass values between the two states under consideration. A difference of ± 0.5 Da was considered a significance threshold for deuterium exchange [64]. Deuteros 2.0 [65] was used to generate coverage maps and Woods plots with peptide level significance testing.

### Synthesis of fluorescein labeled 11-DOC

Unless noted, materials and solvents were purchased from MilliporeSigma (Burlington, MA) and used without further purification. FAM-DBCO, 6-isomer was purchased from Lumiprobe Corporation (Hunt Valley, MD) and used without further purification. 5-bromovaleryl chloride was purchased from TCI America (Montgomeryville, PA) and used without further purification. Thin layer chromatography (TLC) was performed on Sorbent Technologies XHL 254 silica gel plates. Premium Rf 60 Å silica gel was used for column chromatography. ^1^H and ^13^C NMR spectra were obtained on a Bruker Avance Neo 400 MHz NMR spectromer or Bruker 500 MHz Avance III HD NMR Spectrometer with deuterated solvent as noted. Electrospray ionization (ESI) mass spectrometry was performed using a Thermo Q Exactive mass spectrometer with a Vanquish liquid chromatography system.

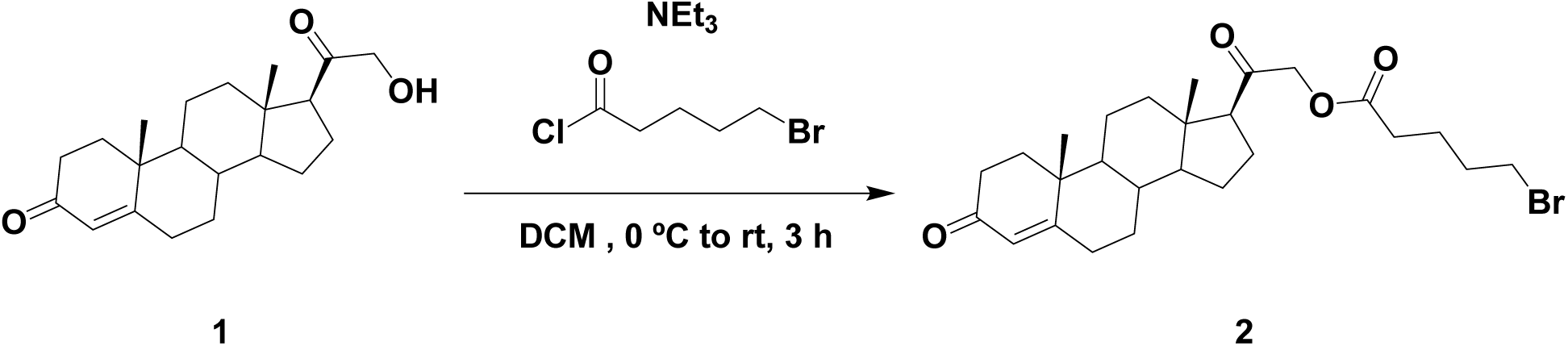

### Synthesis

*2-((10R,13S,17S)-10,13-dimethyl-3-oxo-2,3,6,7,8,9,10,11,12,13,14,15,16,17-tetradecahydro-1H-cyclopenta[a]phenanthren-17-yl)-2-oxoethyl 5-bromopentanoate (****2***). A solution of 21-hydroxyprogesterone (0.186 g, 0.563 mmol, 1.00 eq), triethylamine (0.157 mL, 1.13 mmol, 2.00 eq), and 4 mL dichloromethane (DCM) in a round bottom flask was cooled to 0 °C. To the solution was added 5-bromovaleryl chloride (0.150 mL, 1.20 mmol, 2.12 eq) dropwise over 5 minutes. The solution was allowed to warm to room temperature while continuing to stir for 3 hours. The reaction was diluted with more DCM then washed with water and 1M K_2_CO_3_, dried with MgSO_4_, filtered, and evaporated. Column chromatography was performed (1:1 Ethyl Acetate:Hexanes, Rf = 0.5) to purify the compound, yielding **2** (0.184 g, 0.373 mmol, 66 %) as a white solid. ^1^H NMR (400 MHz, CDCl_3_) δ 5.73 (1H, s), 4.74 (1H, d, J = 17), 4.51 (1H, d, J = 17), 3.42 (2H, t, J = 6.5), 2.52-0.92 (complex, 26H), 1.17 (3H, s), 0.69 (3H, s); ^13^C NMR (100 MHz, CDCl_3_) δ 203.61, 199.73, 172.58, 171.10, 124.06, 69.22, 59.20, 56.29, 53.67, 50.92, 44.79, 38.69, 38.44, 35.80, 35.65, 34.02, 33.24, 32.86, 31.98, 31.90, 24.58, 23.50, 22.96, 21.10, 17.46, 13.30.

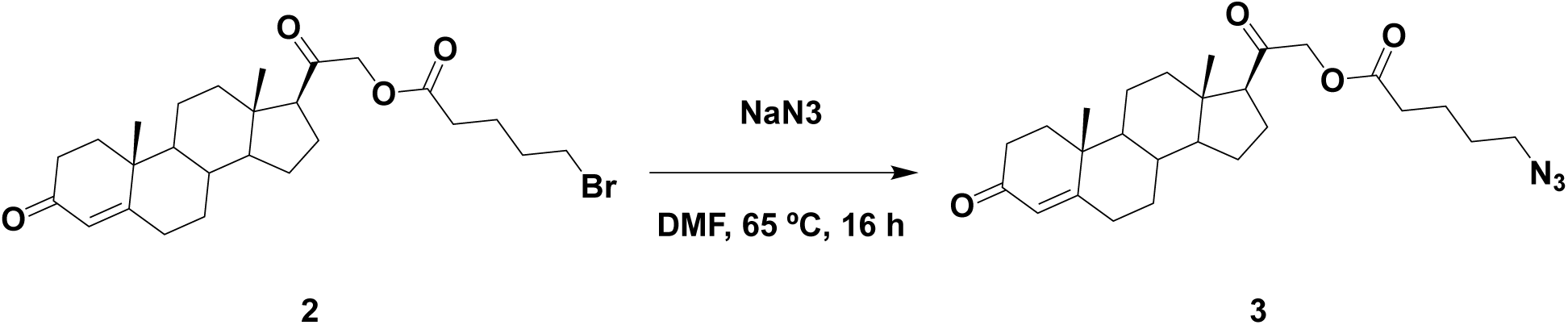

*2-((10R,13S,17S)-10,13-dimethyl-3-oxo-2,3,6,7,8,9,10,11,12,13,14,15,16,17-tetradecahydro-1H-cyclopenta[a]phenanthren-17-yl)-2-oxoethyl 5-azidopentanoate (****3***). To a solution of **2** (0.224 g, 0.454 mmol, 1.00 eq) and 5 mL anhydrous *N,N*-dimethylformamide (DMF) in a round bottom flask, was added sodium azide (0.305 g, 4.69 mmol, 10.3 eq). The solution was was heated to 65 °C for 16 hours. Upon completion, excess sodium azide was filtered off, and the rest was evaporated. The resulting residue was dissolved in Ethyl Acetate and washed three time with water. The organic phase was dried with MgSO_4_, filtered, and evaporated. Column chromatography was performed (4:3 Ethyl Acetate:Hexanes, Rf = 0.57) to purify the compound, yielding **3** (0.162 g, 0.356 mmol, 78 %) as a white solid. ^1^H NMR (500 MHz, CD_3_CN) δ 5.66 (1H, s), 4.76 (1H, d, J = 17), 4.57 (1H, d, J = 17), 3.35 (2H, t, J = 6.5), 2.63-0.98 (complex, 26H), 1.21 (3H, s), 0.68 (3H, s); ^13^C NMR (125 MHz, CD_3_CN) δ 204.85, 199.46, 173.34, 172.11, 124.09, 70.09, 59.55, 56.82, 54.48, 51.69, 45.23, 39.41, 38.88, 36.47, 36.24, 34.55, 33.63, 33.22, 32.78, 28.71, 25.04, 23.34, 22.80, 21.73, 17.61, 13.44.

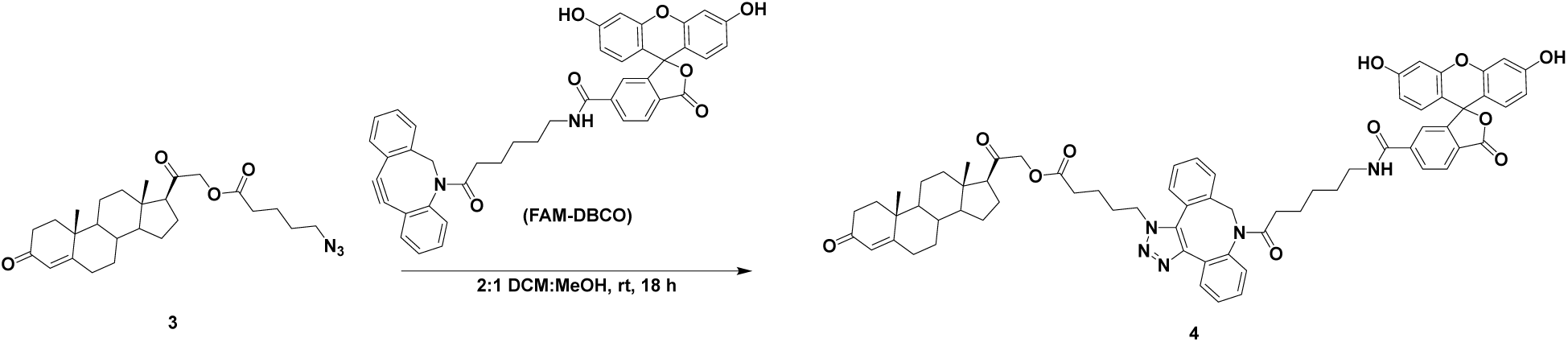

*2-((10R,13S,17S)-10,13-dimethyl-3-oxo-2,3,6,7,8,9,10,11,12,13,14,15,16,17-tetradecahydro-1H-cyclopenta[a]phenanthren-17-yl)-2-oxoethyl 5-(8-(6-(3’,6’-dihydroxy-3-oxo-3H-spiro[isobenzofuran-1,9’-xanthene]-6-carboxamido)hexanoyl)-8,9-dihydro-1H-dibenzo[b,f][1,2,3]triazolo[4,5-d]azocin-1-yl)pentanoate (****4***). A solution of **3** (0.027 g, 59.3 μmol, 1.22 eq), **FAM-DBCO** (0.033 g, 48.8 μmol, 1.00 eq), 6 mL DCM, and 3 mL methanol were combined in a round bottom flask and stirred at room temperature for 18 hours. The solution was evaporated and the compound was purified by reverse phase preparative HPLC (Agilent Technologies) using a ramp of 0% to 100% B over 10 minutes (retention time: 6.18 minutes) to yield **4** (0.009 g, 7.95 μmol, 13 %) as a yellow solid. ESI-MS *m/z* [M + H]^+^ observed: 1132.5, calculated: 1132.5.

### Ensemblator Analysis

For structural comparison of WT and mutant ensembles, 100 conformations were obtained from each trajectory, based on combined clustering results. Structures selected represent the lowest RMSD members of the most populated WT or mutant clusters. Subgroups were identified as WT versus mutant groups. Briefly, Ensemblator performs local conformation comparisons by calculating a local overlaid dipeptide residual (LODR) score to measure residue-level backbone similarity [46, 47]. Global comparisons are performed following a least-squares overlay of all structures using common atoms. From both local and global comparisons, a discrimination index is calculated to access the significance of differences for each atom in both groups. Inter-subgroup variations are calculated as well as intra-subgroup comparisons. The DI is calculated for each atom as the mean of the pairwise distances between the groups minus the mean of the pairwise distances within the group, divided by the higher of the two values. Values range between 0 and 1, going from indistinguishable to structurally distinct ensembles.

## Acknowledgements

The authors wish to thank Dr. Suzanne G. Mays for critical reading and insightful comments on this manuscript. We also thank Dr. Andrew Karplus for in-depth discussion and guidance with the Ensemblator tool.

## Supplemental Information

**Table S1:**
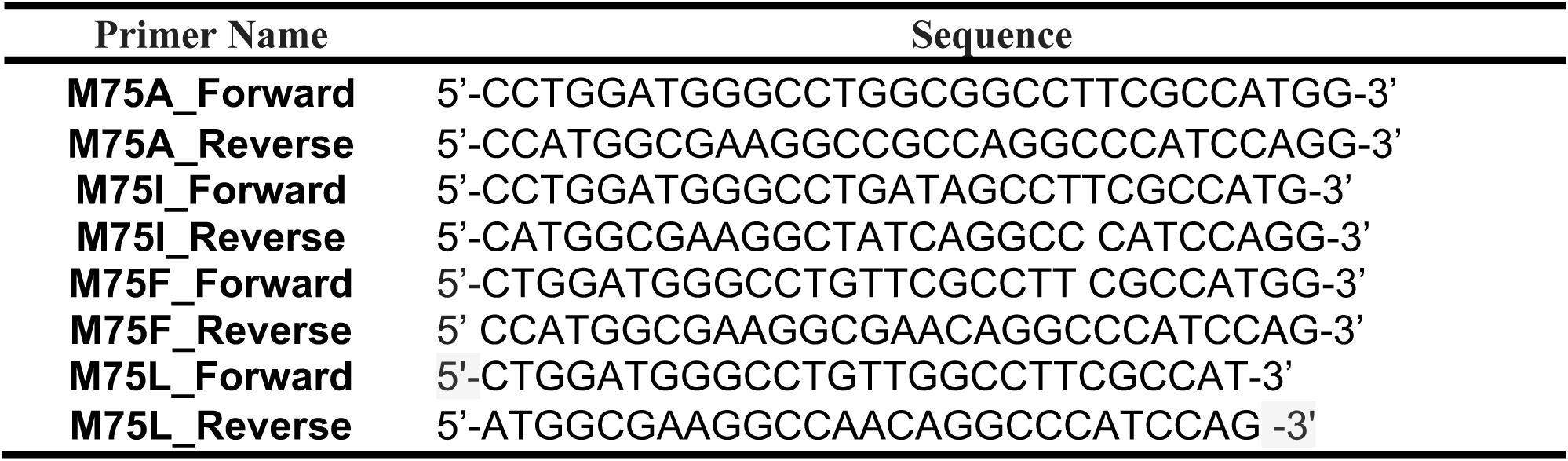
Oligonucleotide primers used for site-directed mutagenesis.

**Figure S1.**
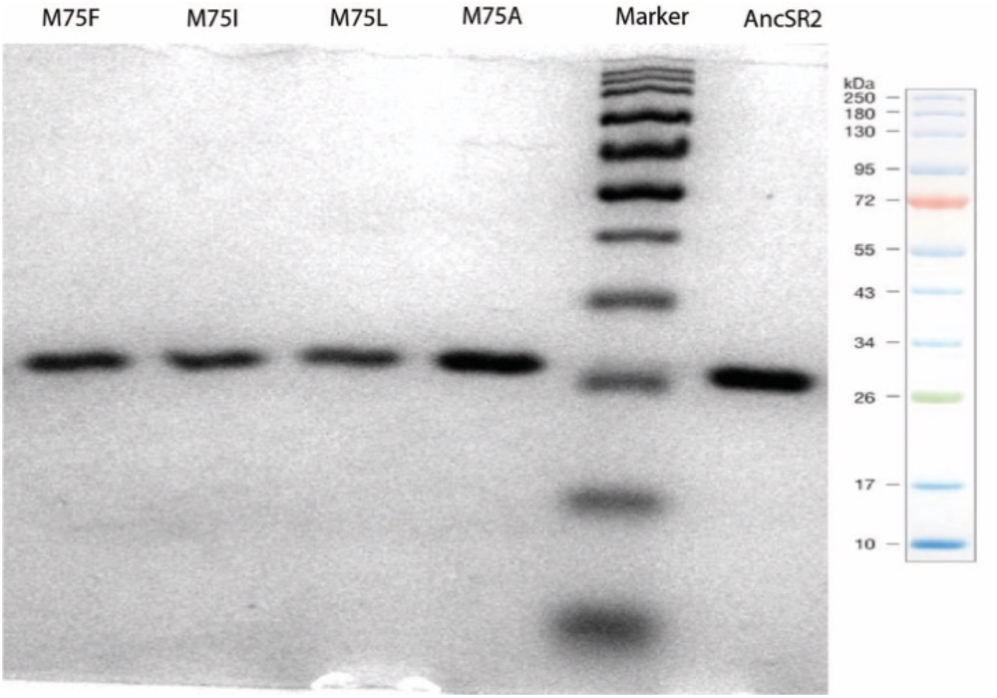
Purity of the LBDs were assessed on 14% SDS-PAGE.

**Figure S2.**
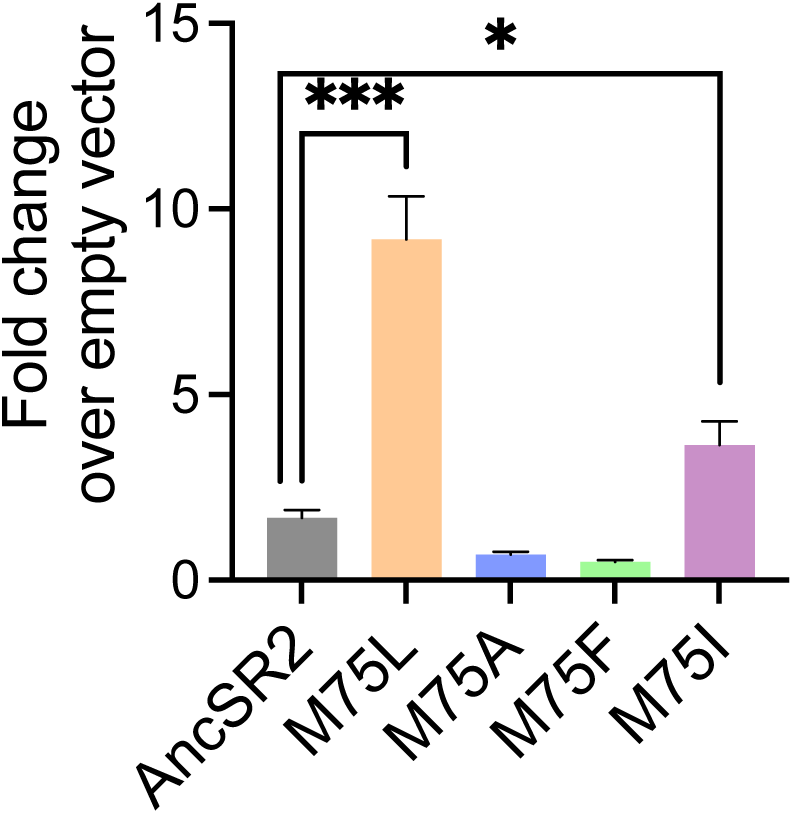
Fold change of AncSR2 and variants over empty vector in the absence of ligand, using CHO cells, showing that constitutive activity of the M75L receptor was independent of the cell line used. Error bars associated with each column represent SEM from three independent replicates. Two-tailed unpaired t-test, (***) P < 0.0002, (*) P < 0.05.

**Figure S3.**
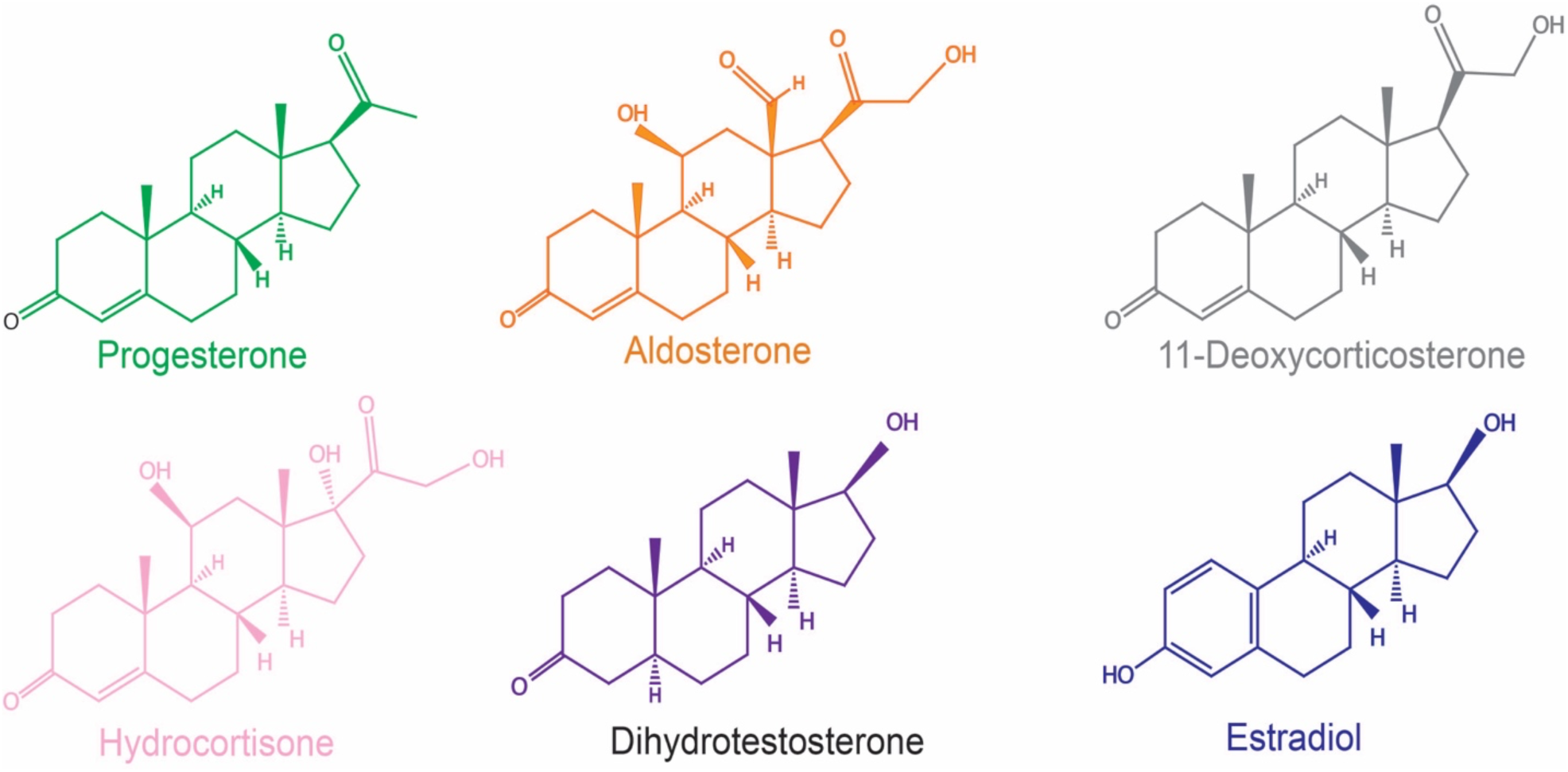
Chemical structures of hormones used in the study.

**Figure S4.**
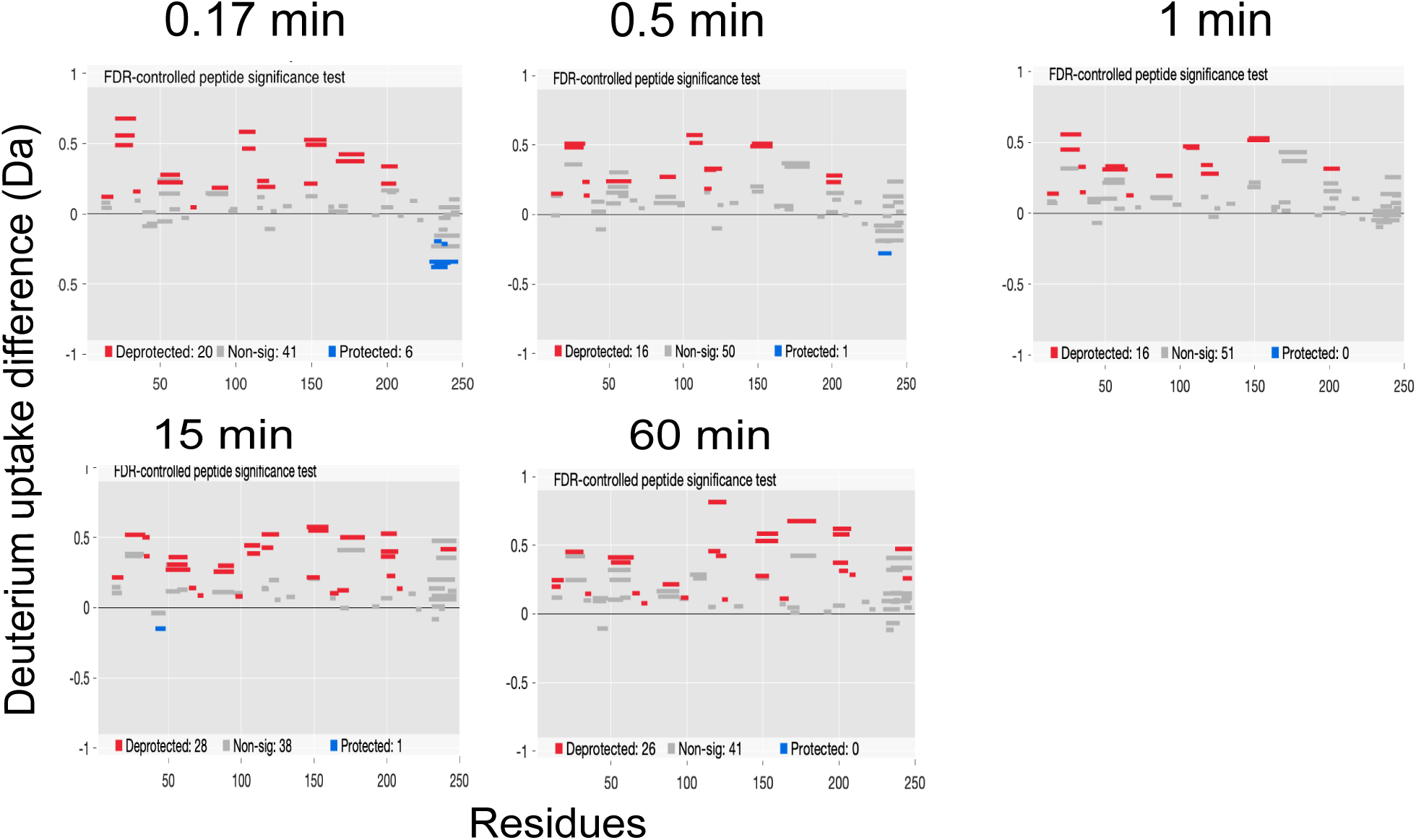
Deuterium exchange difference plot (Woods plot) between M75L and WT AncSR2 apo states (M75LApo – WTApo) at multiple time points. A positive difference indicates increased exchange in apo M75L while a negative difference represents decrease in the deuterium exchange.

**Figure S5:**
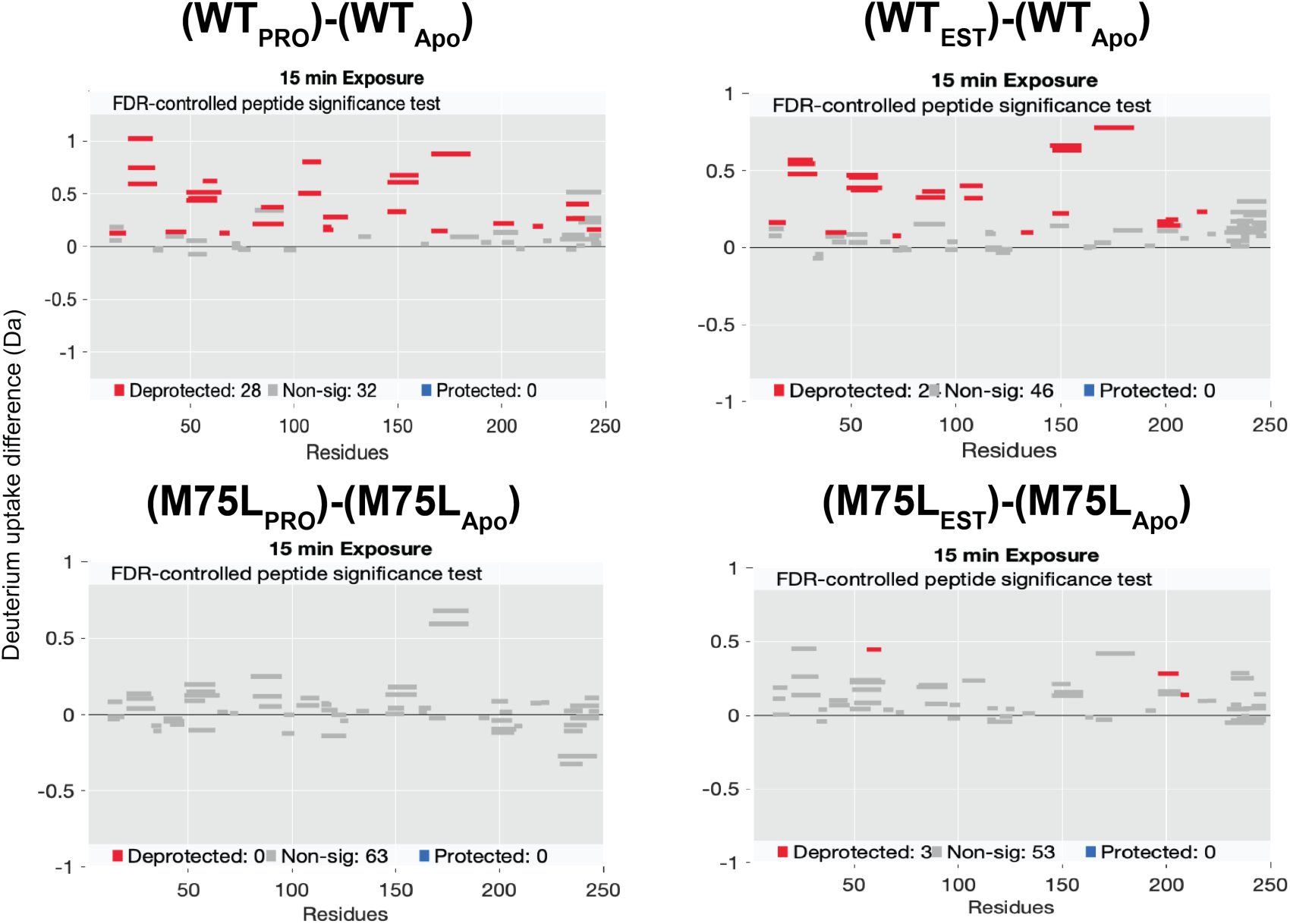
Deuterium exchange difference plots (Time = 15 min) between M75L and AncSR2 apo and ligand bound states. AncSR2 in the PRO and EST bound states showed significantly increased deuterium exchange. In M75L-PRO bound states, no uptake difference was observed in most protein regions, while increased exchange (close to 0.5 Da significance threshold) was seen in the 55-63 peptide of M75L-EST.

**Figure S6:**
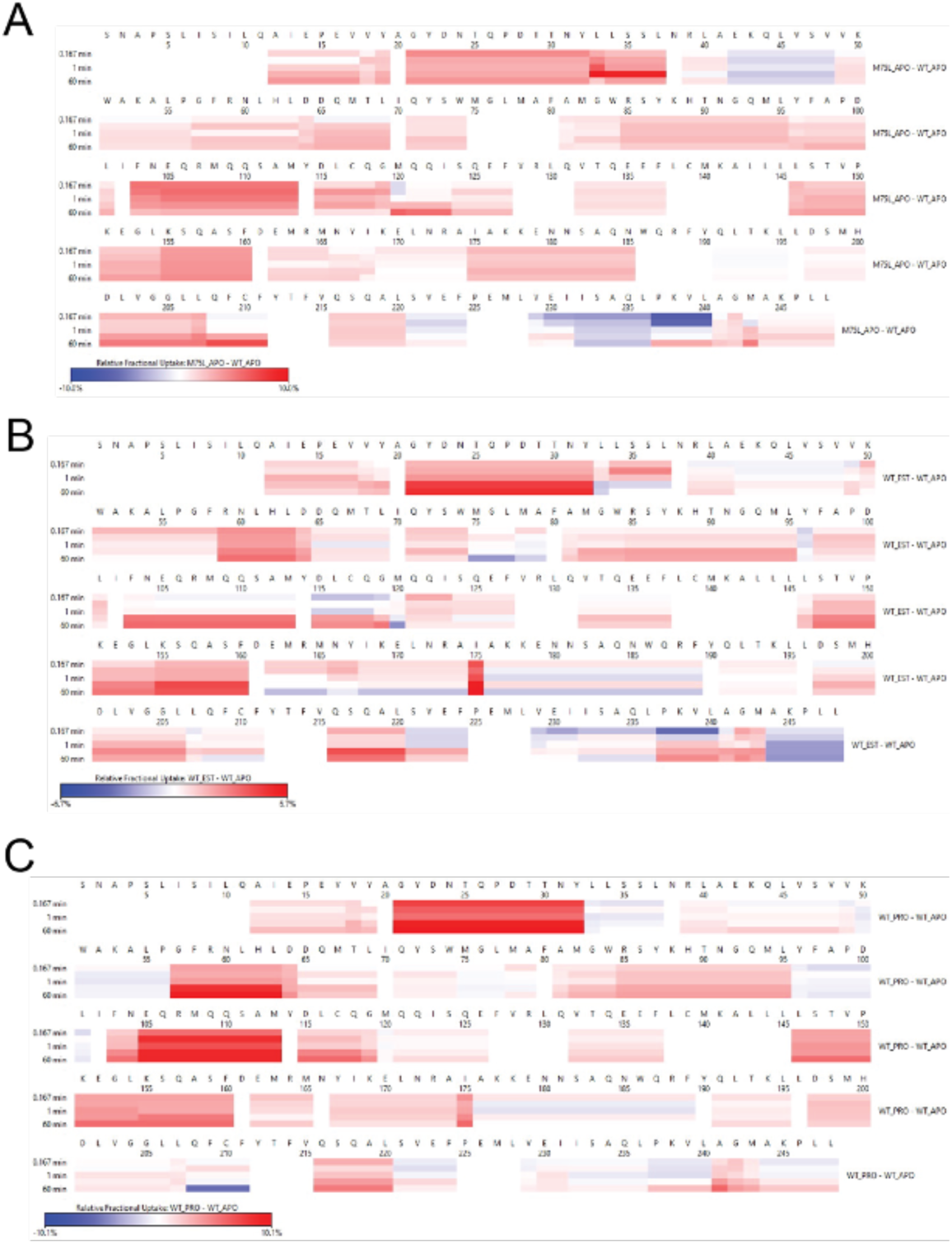
Difference in relative fractional uptake between A) apo states of M75L and WT AncSR2, B) WT AncSR2 apo and estradiol bound (WT-EST) states and C) WT AncSR2 apo and progesterone bound (WT-PRO) states.

**Figure S7:**
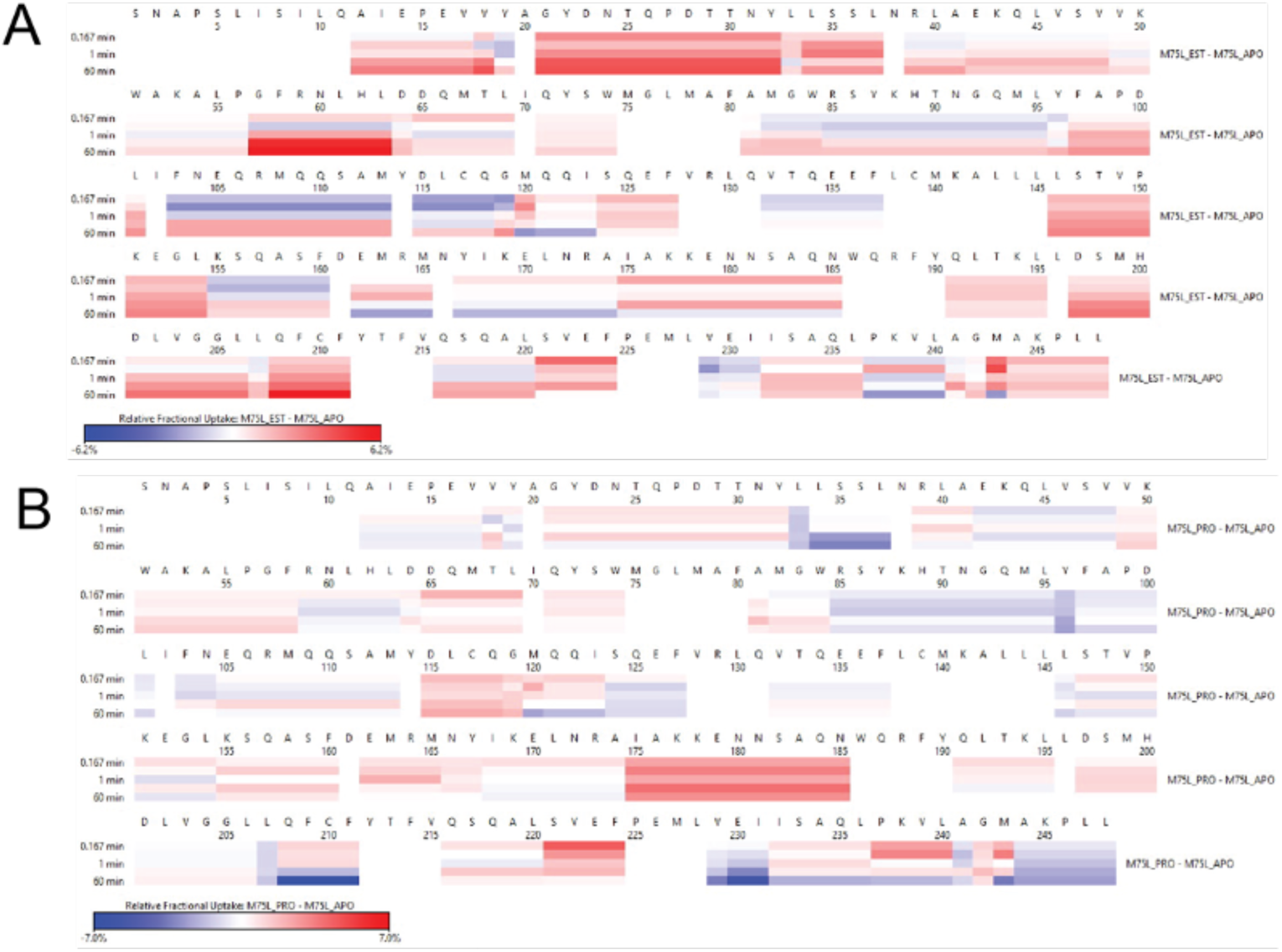
Difference in relative fractional uptake between A) M75L apo and estradiol-bound (M75L-EST) states, B) M75L apo and progesterone-bound (M75L-PRO) states.

**Figure S8.**
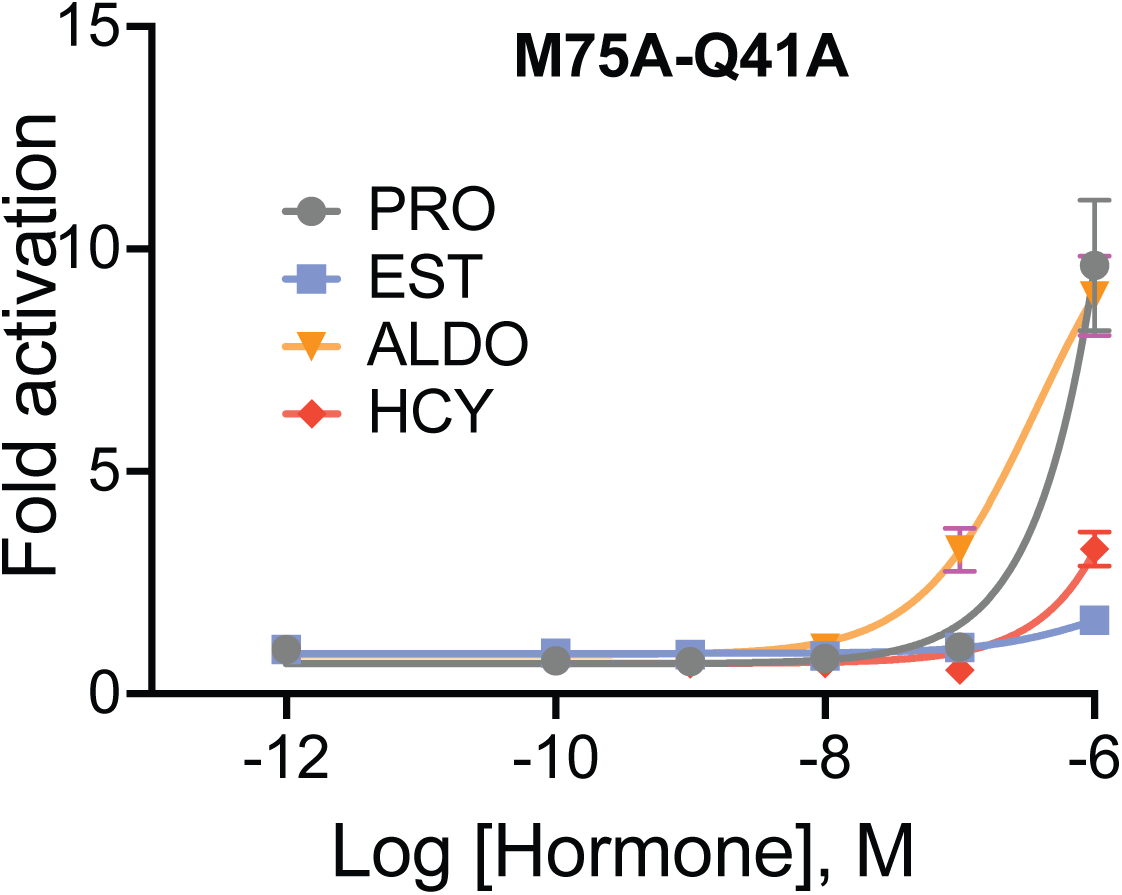
Transcriptional activation in the M75A mutant is abolished upon substitution of Q41A (I.e. in the M75A-Q41A double mutant). Assay was performed with four ligands: progesterone (PRO), estradiol (EST), aldosterone (ALDO) and hydrocortisone (HCY). Each data point is an average of six independent replicates from two biological experiments. Error bars associated with each data point represent SEM.

**Figure S9.**
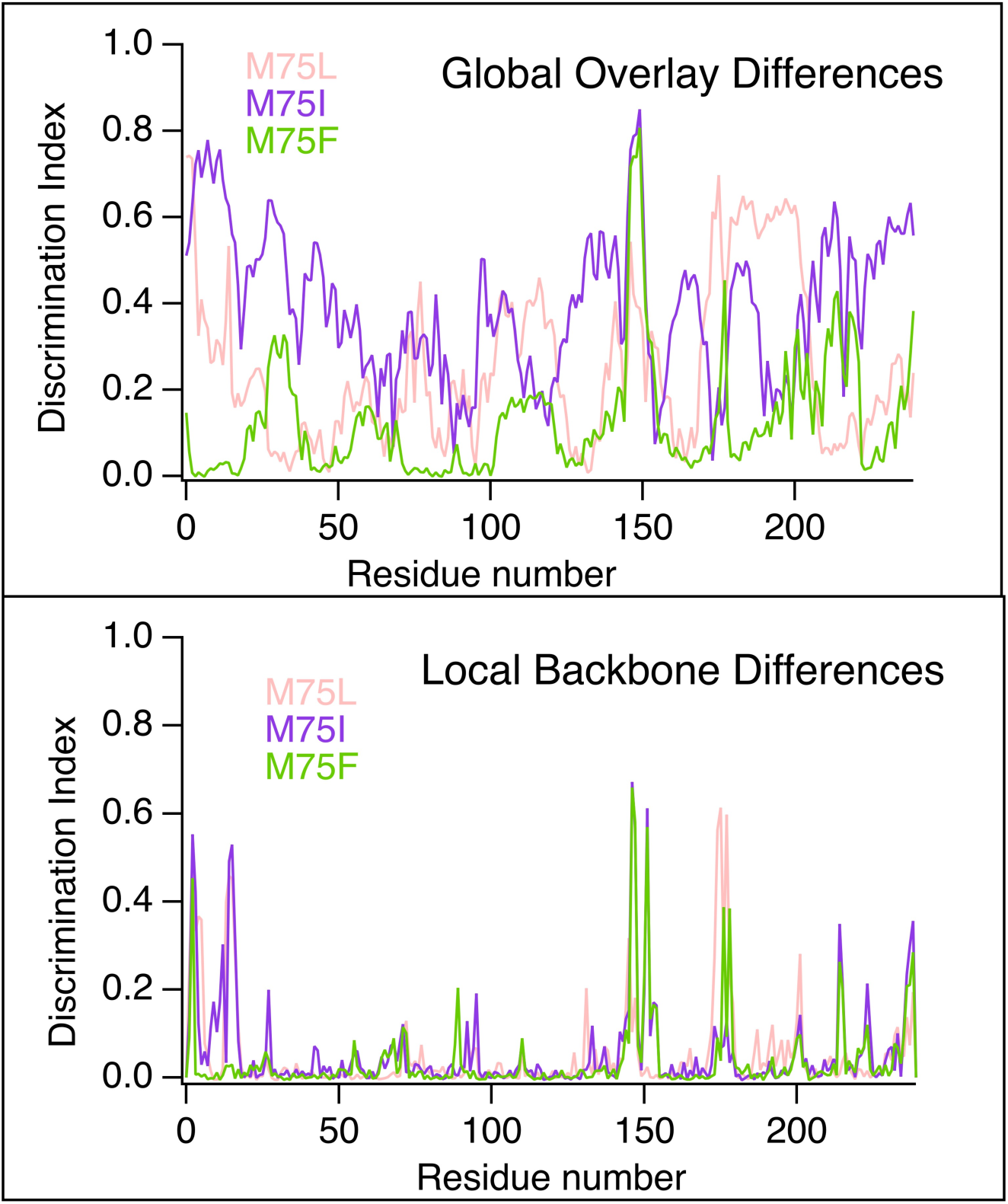
Discrimination Scores reporting on the global and local conformational differences between ensembles of WT AncSR2 and mutants.

